# Relish plays a dynamic role in the niche to modulate *Drosophila* blood progenitor homeostasis in development and infection

**DOI:** 10.1101/2021.04.04.438424

**Authors:** Parvathy Ramesh, Nidhi Sharma Dey, Aditya Kanwal, Sudip Mandal, Lolitika Mandal

## Abstract

Immune challenges demand the gearing up of basal hematopoiesis to combat infection. Little is known about how during development, this switch is achieved to take care of the insult. Here, we show that the hematopoietic niche of the larval lymph gland of *Drosophila* senses immune challenge and reacts to it quickly through the nuclear factor-κB (NF-κB), Relish, a component of the immune deficiency (Imd) pathway. During development, Relish is triggered by ecdysone signaling in the hematopoietic niche to maintain the blood progenitors. Loss of Relish causes an alteration in the cytoskeletal architecture of the niche cells in a Jun Kinase dependent manner, resulting in the trapping of Hh implicated in progenitor maintenance. Notably, during infection, downregulation of Relish in the niche tilts the maintenance program towards precocious differentiation, thereby bolstering the cellular arm of the immune response.

## Introduction

The larval blood-forming organ, lymph gland, is the site for definitive hematopoiesis in *Drosophila* (Banerjee et al., 2019; Evans et al., 2003; Jung, 2005; Lanot et al., 2001; Mandal et al., 2004). Interestingly, there are noticeable similarities between the molecular mechanisms that regulate the lymph gland and those essential for progenitor-based hematopoiesis in vertebrates (Evans et al., 2003; Gold & Brückner, 2014). The lymph gland is formed in embryonic stages, and through various larval stages, it grows in size. The mature third instar larval lymph gland is a multi-lobed structure with well-characterized anterior lobe/primary lobes that have three distinct zones. The prohemocytes/progenitor cells are medially located and define the medullary zone (MZ), while the differentiated hemocytes populate the peripheral zone or cortical zone of the primary lobe. Posterior to these two zones, a group of cells forms the posterior signaling center (PSC) or the niche. Except for one study that claims otherwise (Billel Benmimoun et al., 2015), several studies demonstrate that PSC/niche maintains the homeostasis of the entire organ by positively regulating the maintenance of these two zones (***Figure 1A*** and ***B***) (Jung, 2005; Kaur et al., 2019; Krzemień et al., 2007; Mandal et al., 2007; Mondal et al., 2011; Sharma et al., 2019). During development, this organ is the site of proliferation, maintenance, and differentiation of hemocytes. Only with the onset of pupation do the lymph glands rupture to disperse the blood cells into circulation (Grigorian et al., 2011).

**Figure 1:**
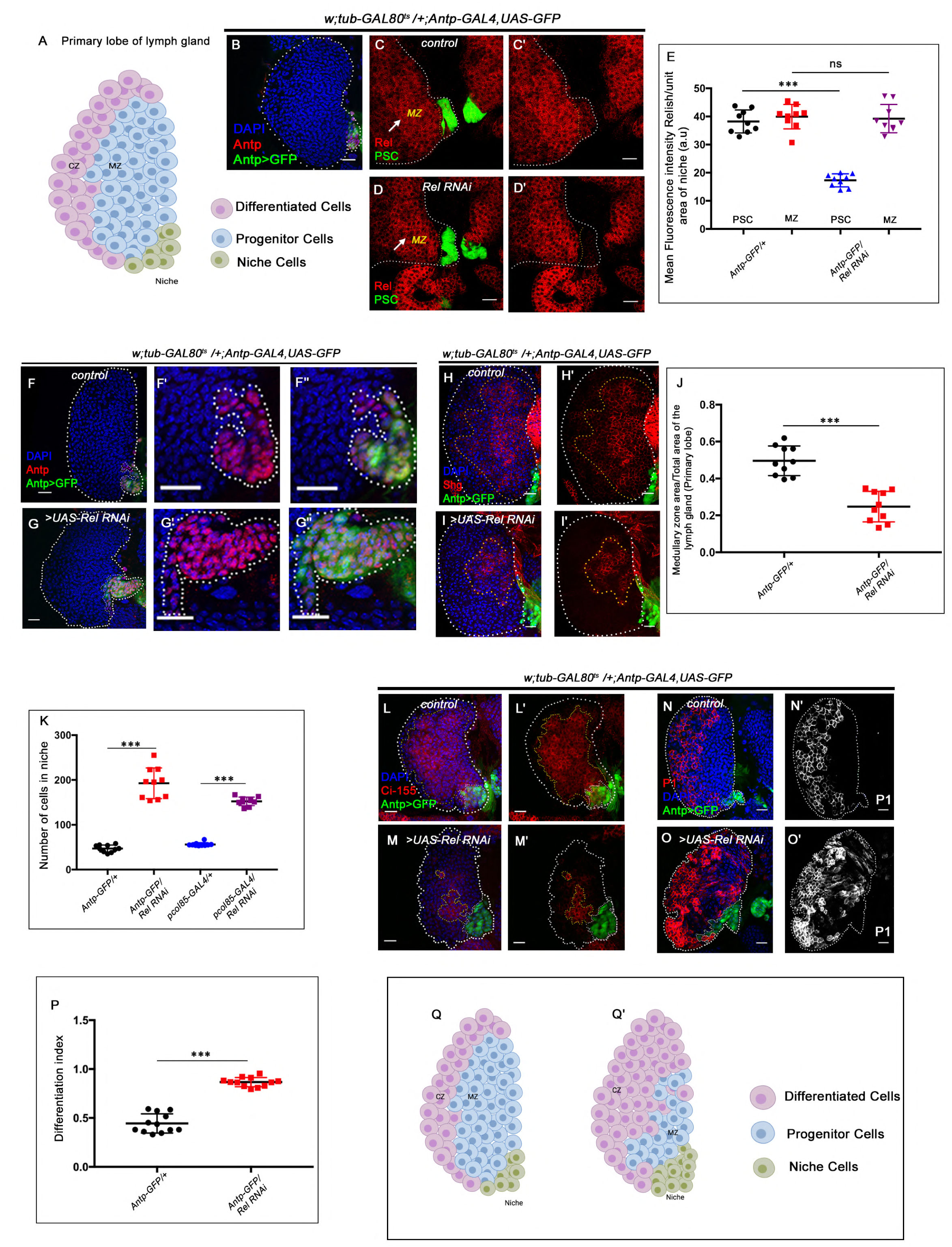
Relish expression and its function in hematopoietic niche of *Drosophila* larval lymph gland. Genotypes are mentioned in relevant panels. Scale bar: 20 µm **A.** Schematic representation of *Drosophila* larval lymph gland with its different cell types. **B.** Hematopoietic niche in larval lymph gland visualized by *Antp-Gal4,UAS-GFP* and Antennapedia (Antp) antibody. **C-D’.** Expression of Relish (Antibody: red) in larval lymph gland. (**C**) Relish is expressed in the hematopoietic niche of lymph gland and in the progenitor population. (**C’**) Zoomed in view of the niche showing the expression of Relish in control niche. (**D-D’**) Relish expression is abrogated in the niche upon RNAi mediated down regulation. **E.** Quantitation of Relish expression in the niche. Significant reduction in Relish expression was observed in niche (n=10, P-value = 7.4×10^-9^, two-tailed unpaired Student’s t-test) whereas progenitor specific expression remained unchanged (n=10, P-value = 0.764, two-tailed unpaired Student’s t-test). **F-G’’.** Effect of Relish loss from the niche on cell proliferation. (**F’-F’’**) Antp expression marks the niche of wild type lymph gland. (**G-G’’**) Loss of Relish function from niche leads to increase in niche cell number. **H-I*’***. Hematopoietic progenitors of larval lymph gland (red, reported by DE-Cadherin (Shg) immunostaining). Compared to control **(H-H’)**, drastic reduction in progenitor pool was observed when Relish function was attenuated from niche (**I-I’**). **J.** Quantitation of Shg positive progenitors population Relish knockdown from the niche using *Antp-GAL4* (n=10, P-value =8.47×10^-6^, two-tailed unpaired Student’s t-test) **K.** Quantitation of niche cell number upon Relish knockdown from the niche using *Antp-GAL4* (n=10, P-value =1.3×10^-7^, two-tailed unpaired Student’s t-test) and *pcol85-GAL4* (n=11, P-value =1.2×10^-12^, two-tailed unpaired Student’s t-test). **L-M’.** Hematopoietic progenitors of larval lymph gland (red, reported by Ci^155^immunostaining) (**L-L’**). Loss of Relish from the niche resulted in reduction in Ci^155^ positive progenitor pool (**M-M’**). **N-O’.** Compared to control (**N-N’**) increase in the amount of differentiated cell population (red, P1 immunostaining) was observed upon niche specific downregulation of Relish (**O-O’**). **P**. Quantitative analysis of (**N-O’**) reveals significant increase in the amount of differentiated cells in comparison to control (n=10, P-value =2.3×10^-9^, two-tailed unpaired Student’s t-test). **Q-Q’.** Scheme based on our observation. The white dotted line mark whole of the lymph gland in all cases. In all panels age of the larvae is 96 hrs AEH. The nuclei are marked with DAPI (Blue). Error Bar: Standard Deviation (S.D). Individual dots represent biological replicates. Error Bar: Standard Deviation (S.D). Data are mean±s.d. *P<0.05, **P<0.01 and ***P<0.001.

It is fascinating to note how this reserve population within the lymph gland is prevented from precociously responding to all of the environmental challenges during the entire larval life. Interestingly, during infection, the lymph gland releases the differentiated hemocytes into circulation in larval stages (Khadilkar et al., 2017; Sorrentino et al., 2002).

The three *Drosophila* NF-κB factors, Dorsal, Dorsal-related immunity factor (DIF), and Relish regulate the insect humoral immunity pathway that gets activated during infection (Govind, 1999; Hetru & Hoffmann, 2009). *Drosophila* NF-κB signaling pathways show conspicuous similarity with vertebrates. NF-κB family consists of five members - RelA (p65), RelB, c-Rel, p50/p105, and p52/p100 (Ganesan et al., 2010). In vertebrates, these factors are critical for producing cytokines, regulating cell death, and controlling cell cycle progression (Gilmore, 2006). In *Drosophila*, Dorsal and Dif activation happens during embryogenesis as well as during gram-positive bacterial and fungal infections. In both cases, it is triggered by the activation of the Toll pathway by cleaved cytokine Spatzle (Valanne et al., 2011).

On the other hand, gram-negative bacterial infections activate the Imd pathway. The diaminopimelic acid (DAP)-type peptidoglycan from the cell wall of the bacteria directly binds to the peptidoglycan recognition protein-LC (Choe, 2002; Gottar et al., 2002; Kaneko et al., 2006; Rämet et al., 2002) or peptidoglycan recognition protein-LE {PGRP-LC or PGRP-LE}. This binding initiates a signaling cascade that elicits the cleavage, activation, and nuclear translocation of Relish with the subsequent transcription of antimicrobial peptide genes (Choe, 2002; Hedengren et al., 1999).

IMD pathway has been studied intensively in the context of immunity and inflammation, but far less is understood about the developmental function of this pathway. Accumulating evidence from studies, however, suggests that the IMD pathway may also have distinct roles in development. In *Drosophila,* Relish and its target genes are activated during neurodegeneration and overexpression of Relish during development causes apoptosis in wing disc cells, neurons, photoreceptors (Cao et al., 2013; Chinchore et al., 2012; Katzenberger et al., 2013; Tavignot et al., 2017) and autophagy in salivary gland cell (Nandy et al., 2018). These studies point out to diverse developmental requirements of Relish beyond immunity in *Drosophila*. Since IMD is an evolutionarily conserved signaling cascade, *Drosophila,* therefore, turns out to be a great model to explore diverse function of the components of this pathway.

Expression of Relish in the hematopoietic niche of the lymph gland during non-infectious conditions prompted us to investigate its role in developmental hematopoiesis. We found that Relish acts as an inhibitor of c-Jun Kinase Signaling *(JNK)* in the hematopoietic niche. During infection, Relish is known to inhibit JNK signaling through *tak1* in *Drosophila* (Park, 2004). Interestingly, we found similar crosstalk being adopted during development in the hematopoietic niche. Activation of JNK signaling in *Drosophila* is associated with alteration of the cytoskeletal architecture of cells during various developmental scenarios, including cell migration, dorsal closure, etc (Homsy et al., 2006; Jacinto et al., 2000; Kaltschmidt et al., 2002; Kockel et al., 2001; Rudrapatna et al., 2014). We found that upon Relish loss, JNK activation causes upregulation of actin remodelers, Enabled and Singed in the niche. The actin cytoskeletal remodeling, in turn, affects the formation of cytoneme-like filopodial projections that help in transporting Hh from the niche to the adjoining progenitors (Mandal et al., 2007; Pennetier et al., 2012; Y. Tokusumi et al., 2012). As a consequence, proper communication between niche and progenitor cells gets disrupted, leading to precocious differentiation at the expense of progenitors.

The hematopoietic niche maintains the delicate balance between the number of progenitors and differentiated cells of the lymph gland (Krzemień et al., 2007; Mandal et al., 2007; Mondal et al., 2011). During development, this organ accumulates hemocytes for post-larval requirements. However, during infection, this organ precociously releases the content into circulation (Lanot et al., 2001). Therefore, a switch is essential to enable the transition from basal hematopoiesis towards the emergency mode to enable the organism to combat infection. The pathway identified in this study, critical for niche maintenance and developmental hematopoiesis, is also exploited during the immune challenge. The circuit engaged in niche maintenance and, therefore, crucial for developmental hematopoiesis gets disrupted during infection. We found that Relish in the niche serves as a joystick to achieve control between developmental and immune response.

Previous studies have demonstrated that Relish needs to be activated in the fat body to mount an immune response (Cha et al., 2003; Charroux & Royet, 2010). We show that to reinforce the cellular arm of the innate immune response, Relish needs to be downregulated in the niche during infection. Though the candidate that breaks the maintenance circuit remains to be identified, nonetheless, our study illustrates that the hematopoietic niche can sense the physiological state of an animal to facilitate a transition from normal to emergency hematopoiesis.

## Results

### The hematopoietic niche requires Relish during development

*Drosophila* NF-κB like factor Relish has been studied extensively as a major contributor of humoral immune defense mechanism against gram-negative bacterial infections (Buchon et al., 2014; Ferrandon et al., 2007; Ganesan et al., 2010; Gottar et al., 2002; Kleino & Silverman, 2014). During larval development, Relish expresses in the hematopoietic niche {marked by *Antp-GAL4>UAS-GFP*, a validated reporter for niche cells (***Figure 1C-C’***)}. In addition to the niche, the hemocyte progenitor cells (MZ) also express Relish (arrow, MZ, ***Figure 1C***). The niche-specific expression was further validated by the down-regulation of Relish using *UAS-Relish RNAi* within the niche that resulted in complete loss of Relish protein therein (***Figure 1D-D’***). As evident from the quantitative analysis (***Figure 1E***) of the above data, the expression of Rel in the niche was drastically affected while that of the MZ is comparable to the control. Whether this transcription factor executes any role in developmental hematopoiesis, beyond its known role in immune response, inspired us to carry out *in vivo* genetic analysis using *Drosophila* larval lymph gland.

We employed the TARGET system (McGuire et al., 2004) to investigate the role of Relish, if any, in the hematopoietic niche. Compared to the control, wherein the number of cells in the hematopoietic niche ranges from 40-45 (***Figure 1F-F’’***), a niche-specific down-regulation of Relish results in a four-fold increase in the cell number (***Figure 1G-G’’*** and ***K***). A similar increase is evidenced upon down-regulation of Relish by another independent niche-specific driver, *collier-GAL4* (Krzemień et al., 2007) (***Figure 1 figure supplement 1A-B’ and Figure 1K***). To further validate the phenotype, the lymph gland from the classical loss of function of Relish (*Rel^E20^*) was analyzed. Interestingly, compared to control, *Rel^E20^* niches exhibit a two-fold increase in cell number (***Figure 1 figure supplement 1C-D’*** and ***E***). Likewise, overexpression of Relish specifically, in the niche, causes a decline in the cell number (***Figure 1 figure supplement 1F-G’’*** and ***H***).

To investigate if the hyperproliferative niche is still capable of performing its function of progenitor maintenance (Mandal et al., 2007) we assayed the status of the progenitors. Interestingly, compared to the control, the loss of Relish from the niche results in a drastic reduction in the number of the progenitor cells (visualized by DE-Cadherin: Shg (Jung, 2005; Sharma et al., 2019) (***Figure 1H-I’ and J*)** and Cubitus interruptus: Ci^155^ ***Figure 1L-M’***) with a concomitant increment in the number of differentiated hemocytes (visualized by plasmatocyte marker by P1, Nimrod ***Figure 1N-O’***) (Asha et al., 2003; Jung, 2005; Kurucz et al., 2007). Quantitation of differentiation index in the genotype described above reveals a two-fold increase in plasmatocyte number (***Figure 1P***). Moreover, in these lymph glands, the differentiated cells, instead of being spatially restricted in the CZ, are dispersed throughout (***Figure 1N-O’***).

Although the differentiation index increases, there was no induction of lamellocytes (visualized by lamellocyte marker β-PS: myospheriod (Stofanko et al., 2008) : ***Figure 1 figure supplement 1I-J’***). The crystal cell numbers also remains unaltered (***Figure 1 figure supplement 1K-L’*** and ***M***), suggesting a tilt towards plasmatocyte fate upon Relish loss from the niche.

These results collectively indicate that Relish plays a critical role in determining the number of niche cells in the developing lymph gland (***Figure 1Q-Q’***).

### Relish loss from the hematopoietic niche induces proliferation

Our expression analysis throughout development reveals that around 45-48 hrs AEH, Relish can be detected in the niche as well as in the progenitors (***Figure 2 figure supplement 1A-E’***). The co-localization of Rel with validated markers of progenitors like TepIV (Dey et al., 2016; Irving et al., 2005; Kroeger et al., 2012; J. Shim et al., 2013) and Ance (B. Benmimoun et al., 2012) further endorsed Rel’s progenitor specific expression (***Figure 2 figure supplement 1F-F’’***). On other hand, co-labeling with Pxn-YFP, a differentiated cell marker (Nelson et al., 1994), reveals that Rel is downregulated from CZ (***Figure 2 figure supplement 1G-G’***). Therefore, we traced back to post second instar stages to get a better insight into the phenotype that is caused upon Relish loss from the niche. At 54-64 hrs AEH, compared to wild type (***Figure 2A-A’’*** and ***C-C’’***), downregulation of Rel by *Antp-Gal4* results in an increase in EdU incorporation in the niche (***Figure 2B-B’’*** and ***D-D’’)***. In context to the niche, a definite proliferation pattern is observable during development. Compared to the rest of the lymph gland, niche cell proliferation decreases by 86 hrs AEH (***Figure 2E-E’’*** and ***2I***). Beyond this time point, EdU incorporation rarely occurs in the niche (***Figure 2G-G’’*** and 2***I***). In sharp contrast to this, upon niche-specific down-regulation of Relish, there is a failure in attaining the steady-state proliferative pattern by 86 hrs AEH. Quite strikingly, EdU incorporation continues even at 96 hrs when the control niche cells have stopped proliferating (Compare ***Figure 2G-G’’*** with ***H-H’’*** and ***Figure 2I***). These proliferating niche cells are indeed mitotically active is evident by the increase in Phospho Histone H3 (PH3) incorporation compared to the control (***Figure 2 J-K’’*** and ***L***). In addition to these snapshot techniques, *in vivo* cell proliferation assay of the niche was done employing the FUCCI system (fluorescent ubiquitination based cell cycle indicator) (Zielke & Edgar, 2015). Fly-FUCCI relies on fluorochrome-tagged probes where the first one is a GFP fused to E2F protein, which is degraded at S phase by Cdt2 (thus GFP marks cells in G2, M, and G1 phase). The second probe is an mRFP tagged to the CycB protein, which undergoes Anaphase

Promoting Complex/cyclosome mediated degradation during mid mitosis (thereby marking cells in S, G2, and M phases). While in control by 96 hrs AEH, niche cells are mostly in G2-M (yellow), and in G1 state (green), in loss of Relish, abundance in S phase can be seen at the expense of G1 (***Figure 2 figure supplement 1H-I’’’’*** and ***J***).

**Figure 2:**
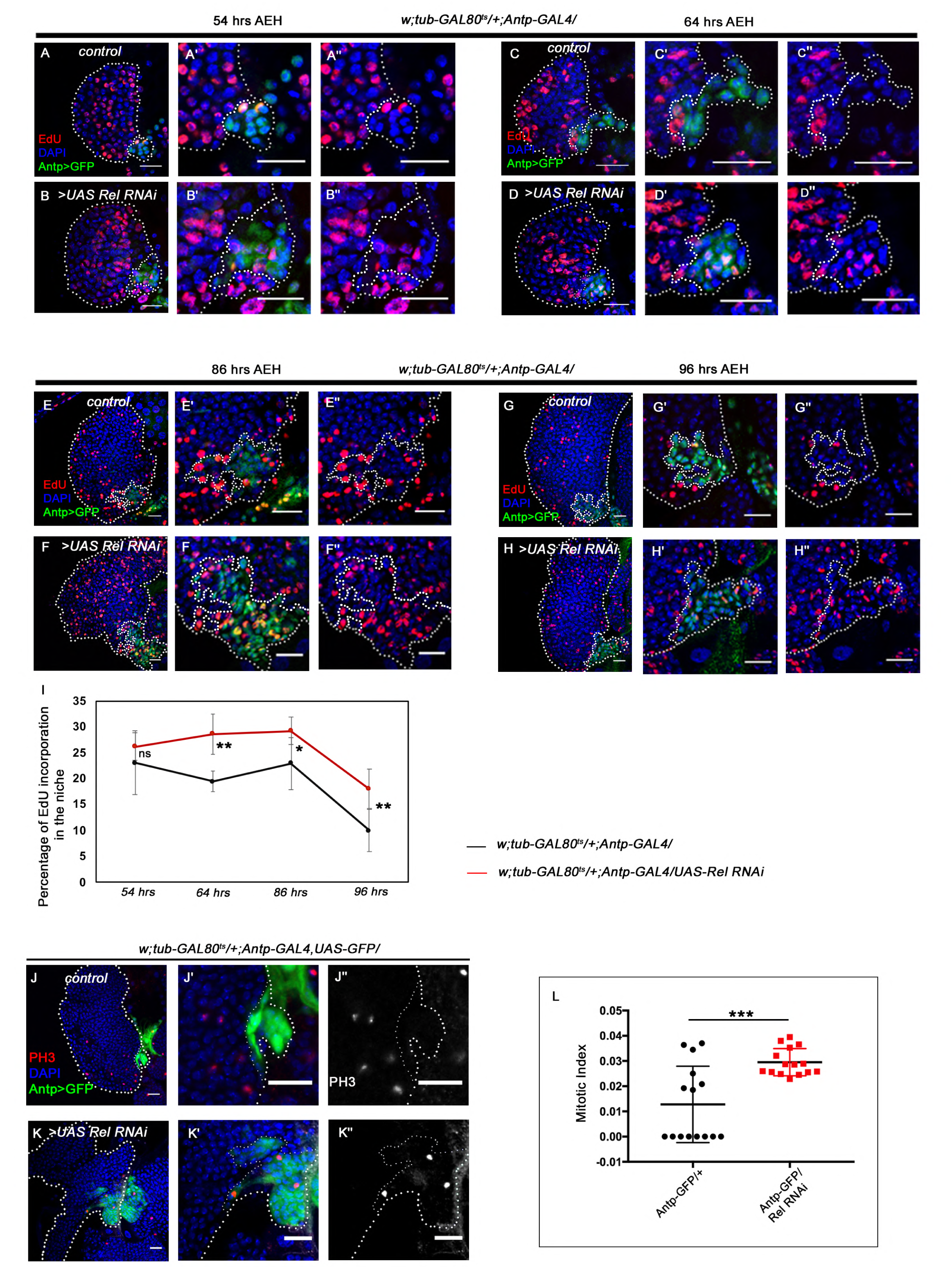
Loss of Relish from the niche causes niche cell hyperplasia. Genotypes are mentioned in relevant panels. Scale bar: 20 µm. Niche is visualized by Antp antibody expression. **A-H’’**. EdU or 5-ethynyl-2’-deoxyuridine marks the cells in S-phase of the cell cycle. EdU profiling at 54hr AEH (**A-B’’**), 64 hrs AEH (**C-D’’**), 86 hrs AEH (**E-F’’**) and 96hr AEH (**G-H’’**) shows EdU incorporation in the niche (green) in control and upon Relish down regulation. Control niches showed scanty EdU incorporation beyond 84 hours (**E-E’’** and **G-G’’**) whereas loss of Relish induced niche cells to proliferate more (**F-F’’** and **H-H’’**). **I.** Graph representing percentage of EdU incorporation in the niche during the course of development in control and Relish loss. Significant increase in the niche cell number is observed with development in Relish loss scenario. (54 hrs, n=6, P-value =.294), (64 hrs, n=6, P-value = 1.3×10^-3^), (86 hrs, n=6, P-value= 2.9×10^-2^), (96 hrs, n=6, P-value = 5.9×10^-3^); two-tailed unpaired Student’s t-test. **J-K’’**. Significant increase in the number of mitotic cells (phospho-histone 3 (PH3), red) was observed upon Relish loss from the niche (**K-K’’**) compared to the control (**J-J’’**). (**L**) Quantitation of the mitotic index of wild type and Relish loss niche (n=15, P-value = 8.1×10^-4^; two-tailed unpaired Student’s t-test). The white dotted line mark whole of the lymph gland in all cases. In all panels age of the larvae is 96 hrs AEH, unless otherwise mentioned. The nuclei are marked with DAPI (Blue). Individual dots represent biological replicates. Error Bar: Standard Deviation (S.D). Data are mean±s.d. *P<0.05, **P<0.01 and ***P<0.001. Error bar: SD

Put together; these results implicate that Relish functions as the negative regulator of niche proliferation in the developing lymph gland.

### Absence of Relish in the niche stimulates proliferation via up regulation of Wingless signaling

Previous studies have shown that the Wingless (Wg) pathway positively regulates niche cell number in addition to its role in the maintenance of the prohemocyte population in the MZ (Sinenko et al., 2009). Upon perturbation of Relish function, a drastic increase in the level of Wingless is evident (arrow, ***Figure 3B-B’’***) in the niche compared to the control (arrow, ***Figure 3A-A’’***). Quantitative analysis reveals a 1.6-fold increase in the fluorescence intensity of Wg per unit area in the niche where Relish’s function is attenuated compared to that of the control (***Figure 3C***). Reducing Wingless function by using a temperature-sensitive mutant allele *wg^ts^* (Bejsovec & Martinez Arias, 1991) following the scheme provided in **Figure 3 figure supplement 1A** led to a decline in niche cell number compared control (Compare ***Figure 3E-E’*** with ***Figure 3D-D’*** and ***H)***, Tweaking of Wg in the background of Rel loss from the niche (***Figure 3F-F’*** and ***H***) restores the hyperproliferative niche (***Figure 3G-G’*** and ***H***), to a cell number comparable to the control (***Figure 3D-D’*** and ***H***).

**Figure 3:**
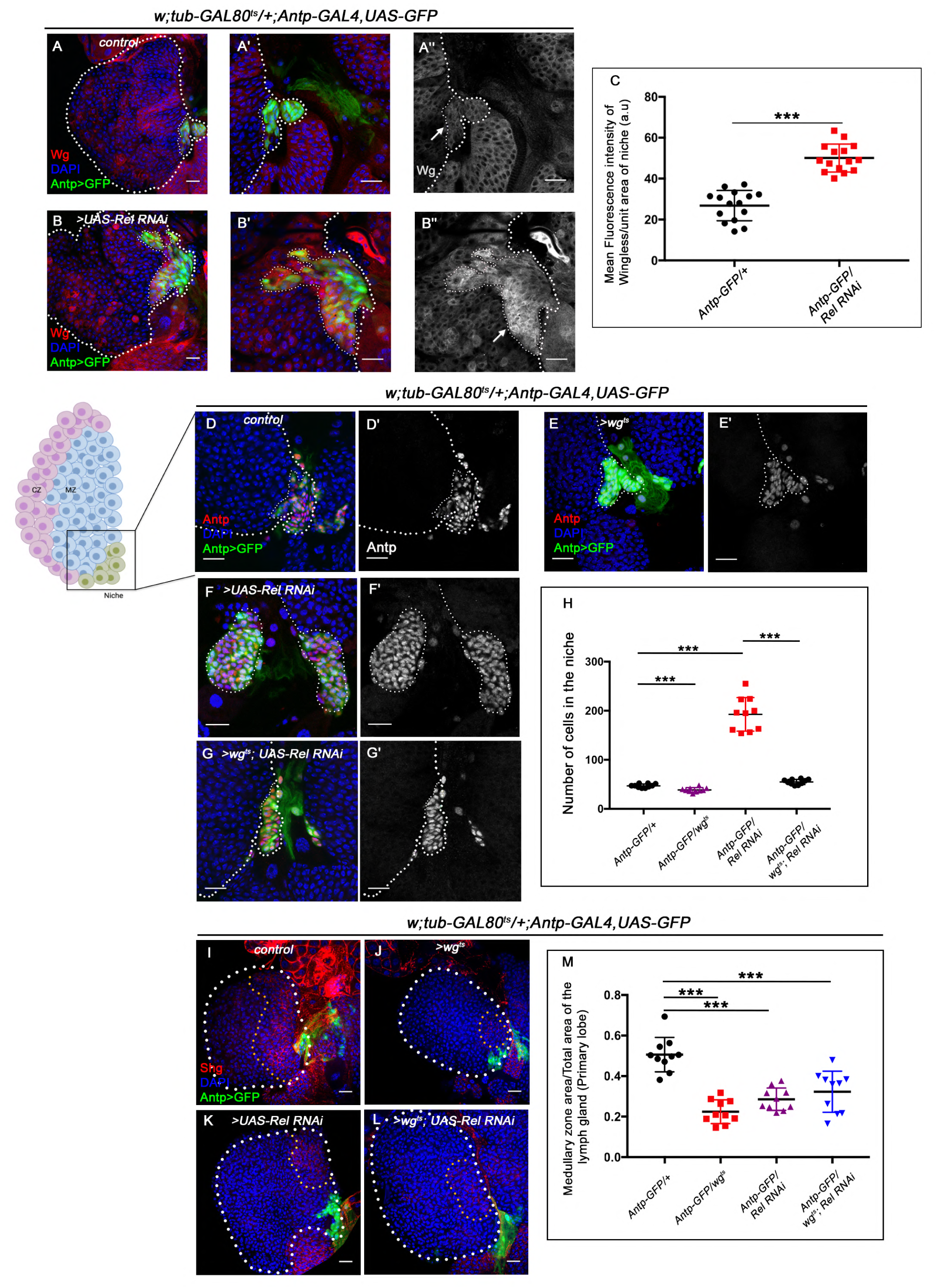
Upregulated Wingless signaling leads to increase in niche number. The genotypes are mentioned in relevant panels. Scale bar: 20µm. **A-B’’**. Expression of Wingless (antibody) in the lymph gland. The hematopoietic niche is visualized by *Antp-GAL4>UASGFP*. (**A’-A’’**) and (**B’-B’’**) are higher magnifications of **A** and **B** respectively. In comparison to the wild type niche (**A-A’’**), Wingless protein levels were substantially high in Relish loss of function (**B-B’’**). (**C**) Statistical analysis reveals elevated wingless expression Relish knockdown PSC (n=15; P-value = 5.8 x10^-9^, two-tailed unpaired Student’s t-test.) **D-G’.** The increased niche number observed upon Relish loss (**F-F’**) is rescued upon reducing Wingless level by the temperature sensitive allele *wg^ts^* (**E-E’**) in Relish loss genetic background **(G-G’**). The rescued niche cell number is comparable to control (**D-D’**). **H.** Statistical analysis of the data in (**D-G’**) (n=10; P-value = 2.4×10^-7^ for control versus Relish RNAi, P-value = 4.3×10^-4^ for control versus *wg^ts^* and P-value = 3.4×10^−7^ for *wg^ts^*; Relish RNAi versus Relish RNAi; two-tailed unpaired Student’s t-test). **I-L.** Hematopoietic progenitors of larval lymph gland (red, reported by DE-Cadherin (Shg) staining immunostaining). Knocking down wingless function using *wg^ts^* resulted in loss of Shg positive progenitors (**J**). Downregulating wg function in Relish loss genetic background was unable to restore the reduction in prohemocyte pool (**L**) observed in Relish loss (**K**) scenario in comparison to control (**I**). **M.** Statistical analysis of the data in (**I-L**) (n=10; P-value = 4.80×10^-6^ for control versus Relish RNAi, P-value = 3.8×10^−4^ for *wg^ts^*; Relish RNAi versus control, P-value = 2.18×10^-7^ for control versus *wg^ts^*; two-tailed unpaired Student’s t-test). The white dotted line mark whole of the lymph gland in all cases. In all panels age of the larvae is 96 hrs AEH. The nuclei are marked with DAPI (Blue). Individual dots represent biological replicates. Error Bar: Standard Deviation (S.D). Data are mean±s.d. *P<0.05, **P<0.01 and ***P<0.001

The above data was further endorsed by co-expression of the RNAi constructs for *wg* and Relish in the niche, which gave similar restoration of the hyperproliferative phenotype (***Figure 3 figure supplement 1J*** and ***K***) that is otherwise seen upon Relish loss (***Figure 3 figure supplement 1H*** and ***K***), which is now similar to that of wild type niches (***Figure 3 figure supplement 1G*** and ***K***). These results collectively implicate that Relish plays a significant role in regulating Wg levels in the niche, thereby controlling niche size. Interestingly, although the niche number is comparable to the control in the above genotypes, the defects in maintenance (***Figure 3I-L*** and ***M, Figure 3 figure supplement 1L-O*** and ***P)***, and differentiation (***Figure 3 figure supplement 1B-E*** and ***F, Figure 3 figure supplement 1Q-T*** and ***U***) observed upon Relish loss from the niche was still evident.

This observation led us to infer that the up-regulated Wg in Relish loss controlled the size and not the function of the niche.

### In absence of Relish, altered cytoskeletal architecture of the niche traps Hh

Various studies have established PSC as the niche for hematopoietic progenitors and have shown that it employs a morphogen Hedgehog for this purpose. It has also been shown that niche expansion correlates to expansion in the progenitor population (Baldeosingh et al., 2018; B. Benmimoun et al., 2012; Mandal et al., 2007; Pennetier et al., 2012; T. Tokusumi et al., 2011). However, in contrast to the above studies, despite a three-fold increment in niche cell number upon Relish down-regulation, we observed a significant reduction in the progenitor pool (***Figure 1L-M’***). Moreover, restoration in the number of niche cells by modulating Wg levels in Relish knockdown condition failed to restore the differentiation defects observed upon downregulating Relish from the niche (***Figure 3 figure supplement 1B-E*** and ***F, Figure 3 figure supplement 1Q-T*** and ***U***). To understand this result, we assayed Hedgehog levels in the niche by using an antibody against Hh protein (Forbes et al., 1993). Interestingly, compared to that of the control, there is a substantial increase in Hh protein in the niche where Relish function is abrogated (***Figure 4A-B’’***). Quantitative analysis reveals an almost two-fold increase in the level of Hh protein in the experimental niche (***Figure 4C***).

**Figure 4:**
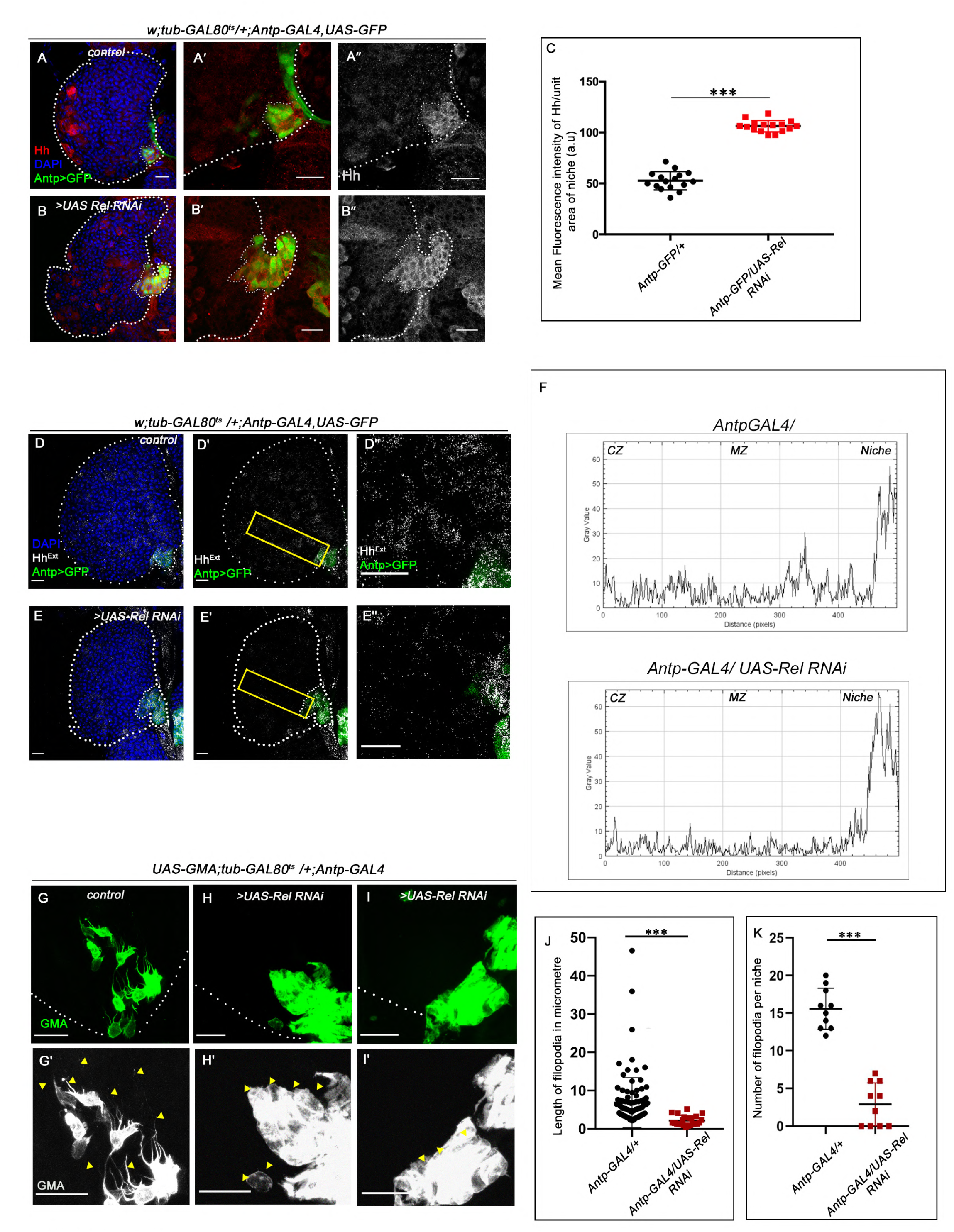
Hedgehog release from the niche is affected in Relish loss of function. The genotypes are mentioned in relevant panels. Scale bar: 20µm. **A-B’’**. Hedgehog (Hh) antibody staining in the lymph gland shows Hh enrichment in the niche. The hematopoietic niche in Relish loss of function (**B-B’’**) exhibits higher level of Hh in comparison to the control (**A-A’’**). **C**. Statistical analysis of fluorescence intensity revealed more than 2.5-fold increase in Hh levels compared to control (n=15, P-value = 2.5×10^-17^, two tailed Students t-test). **D-E’’**. Progenitors of Relish loss of function exhibits lower level of Extracellular Hh (Hh^Extra^) (**E-E’’**) in comparison to those of control (**D-D’’**). **E’’** and **D’’** are zoomed in view of niche and the neighbouring progenitor cells of **E’** and **D’** respectively. The yellow box denotes the area quantified in F. **F.** The intensity profile of Hh^Extra^ in progenitors (along the rectangle drawn from PSC to Cortical zone housing differentiated cells in Figure **4D’** and **E’**) reflects a stark decline in the level of Hh^Extra^ in Relish loss scenario compared to control. **G-I’**. Cellular filopodia emanating from the niche cells were stabilized by using untagged phalloidin. The filopodia in Relish loss of function niches were found to be smaller in length and fewer in number (**H-H’, I-I’**) as compared to control (**G-G’**). **J-K.** Significant reduction in filopodial number (**J,** n=10, P-value = 6.64×10^-9^, two tailed Students t-test) and lengths (**K,** n=6, P-value = 9.19×10^-10^, two tailed Students t-test) were observed in Relish loss scenario compared to control. The white dotted line mark whole of the lymph gland in all cases. In all panels age of the larvae is 96 hrs AEH. The nuclei are marked with DAPI (Blue). Individual dots represent biological replicates. Error Bar: Standard Deviation (S.D). Data are mean±s.d. *P<0.05, **P<0.01 and ***P<0.001.

However, despite having a higher amount of Hh in the niche upon Relish down-regulation, there was a decline in the amount of extracellular Hh (Hh^Ext^) in the prohemocytes compared to control (***Figure 4D-E’’ and F***). This result is in sync with the observation that Rel loss from the niche leads to the reduction in the levels of Ci^155^ in the progenitors (***Figure 1L-M’***), suggesting that Hh produced by the niche, is not sensed by the progenitors resulting in their precocious differentiation.

The alteration in extracellular Hh and decline in Ci^155^ level in the progenitors prompted us to speculate that loss of Relish from niche might have interfered with Hh delivery to the progenitor cells. Several reports in diverse tissues across model organisms have demonstrated filopodia mediated Hh delivery (Bischoff et al., 2013; González-Méndez et al., 2019). Notably, the same holds for the hematopoietic niche cells wherein Hh delivery via filopodia is shown to be relevant (Pennetier et al., 2012; Y. Tokusumi et al., 2012). Therefore, we checked the status of these actin-based cellular extensions emanating from the niche cells in freshly dissected unfixed tissue. For this purpose, *UAS-GMA* (also known as *UAS-moesin-GFP*) that marks F-actin (Kiehart et al., 2000) was expressed in a niche-specific manner. In the control, multiple cellular processes with variable length are detectable, while upon Relish knockdown, filopodial extensions are highly compromised (***arrowheads, Figure 4 G-I’***). Quantitative analyses of the data reveal that both length (***Figure 4J)*** as well as number (***Figure 4 K)*** is altered. As a functional correlate niche specific perturbation of Diaphanous, an actin polymerase resulted in increased differentiation compared to control (***Figure 4 figure supplement 1A-B’*** and ***C)*** Additionally, compared to the control, F-actin (visualized by rhodamine-phalloidin) expression is significantly increased in the cell cortex upon Relish loss from the niche (***Figure 4 figure supplement 2A-B’’*** and ***C***). This accumulation of cortical F-actin intrigued us to further probe into F-actin associated proteins’ status, Singed and Enabled in the niche cells upon loss of Relish. While Singed is the *Drosophila* homolog of Fascin and is involved in cross-linking actin filaments and actin-bundling (Cant et al., 1994; Tilney et al., 2000), Enabled is a cytoskeletal adaptor protein involved in actin polymerization (Gates et al., 2007; T.-Y. Lin et al., 2009). In comparison to the control, where there is a basal level of Singed or lack of Ena expression in the niche, a significant increase in the level of both of these actin-associated proteins occurs upon downregulation of Relish function (Singed: ***Figure 4 figure supplement 2D-E’’*** and ***F*** and Ena: ***Figure 4 figure supplement 2G-H’’*** and ***I***).

These results demonstrate that loss of Relish from the niche induces cytoskeletal rearrangement, which disrupts the proper delivery of Hedgehog to the adjoining progenitors. These results further emphasize how aberrant cytoskeleton architecture might interfere with niche functionality by trapping Hh.

### Ectopic JNK activation leads to precocious differentiation in Relish loss from the niche

Next, we investigated how Relish loss causes alterations in cytoskeletal architecture within the niche. Studies across the taxa have shown Mitogen-activated protein kinases (MAPKs) as a major regulator of cellular cytoskeleton dynamics (Densham et al., 2009; Pichon, 2004; Reszka et al., 1995; Samaj, 2003). The c-Jun-NH2-terminal kinase (JNK) or so-called stress-activated protein kinases, which belong to the MAPK superfamily, are one such key modulator of actin dynamics in a cell. Whether the cytoskeletal remodeling of the niche in the absence of Relish is an outcome of JNK activation was next explored. Compared to the control where there is a negligible level of activation of JNK signaling in the niche, visualized by TRE-GFP: a transcriptional reporter of JNK (Chatterjee & Bohmann, 2012), a robust increase in the expression occurs in the niche where the function of Relish is abrogated (***Figure 5A-B’*** and ***C***). This result implicates that during development, Relish inhibits JNK activation in the hematopoietic niche.

**Figure 5:**
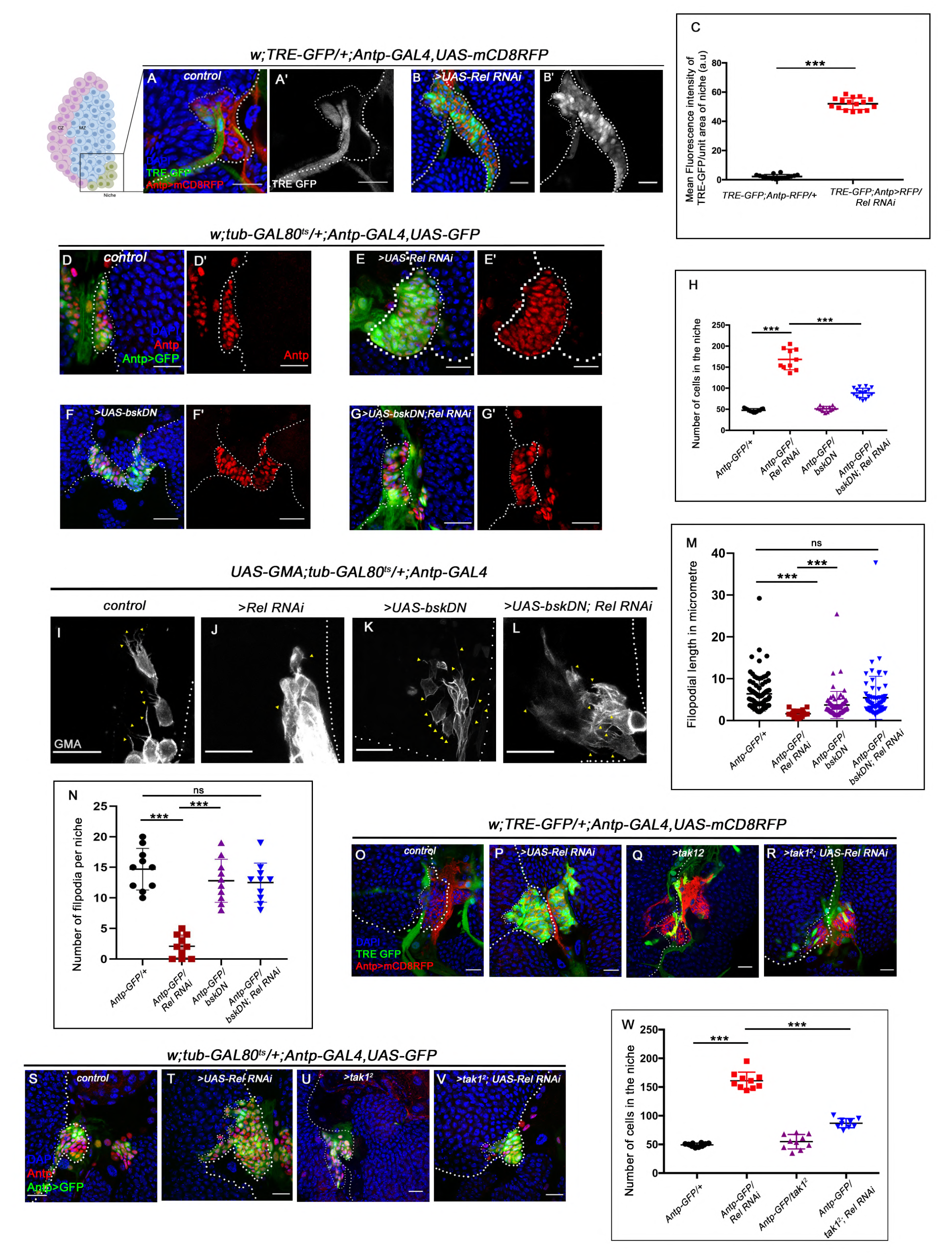
Loss of Relish from the niche activated JNK causing formation of hyperplastic niche. The genotypes are mentioned in relevant panels. Scale bar: 20µm. **A-B’**. Upregulation of JNK signalling visualized by its reporter TRE-GFP (green) in Relish knockdown (**B-B’**) compared with WT niche (**A-A’)**. **C**. Statistical analysis of fluorescence intensity significant increase in TRE-GFP levels compared to control (n=15 P-value = 4.2 x10^-19^, two tailed Students t-test). **D-G’**. Upon simultaneous knockdown of both Relish and JNK from the niche, the increase in niche cell number is observed upon loss of Relish from the niche (**E-E’**) is rescued (**G-G’**) and is comparable to control (**D-D’)** whereas loss of bsk from the niche doesn’t alter niche cell number (**F-F’**). **H**. Statistical analysis of the data in **D-G’** (n=10, P-value = 5.6×10^−8^ for control versus Relish RNAi, P-value = 8.0×10^−7^ for bsk^DN^; Relish RNAi versus Relish RNAi; two-tailed unpaired Student’s t-test). **I-N**. Cellular filopodia from the niche cells in Relish loss of function are found to be smaller in length and fewer in numbers **(J** and **M-N**). Simultaneous loss of both JNK using bsk^DN^ and Relish (**L** and **M-N**) rescued the stunted, scarce filopodia to control state (**I** and **M-N)** whereas loss of JNK did not affect filopodia formation (**K** and **M-N)**. **M-N.** Statistical analysis of the data in **I-L** (Filopodia number: n=10, P=6.96×10^−8^ for control versus Relish RNAi, P-value = 8.11×10^−7^ for bsk^DN^; Relish RNAi versus Relish RNAi, P-value = 0.153 for bsk^DN^ vs control. Filopodia length : n=6, P-value = 2.78×10^-16^ for control versus Relish RNAi, P-value = 1.84×10^-6^ for bsk^DN^; Relish RNAi versus Relish RNAi, P-value = 0.22 for bsk^DN^ vs control; two-tailed unpaired Student’s t-test). **O-R**. Up regulation of JNK signalling visualized by its reporter TRE-GFP (green) in Relish knockdown (**P**) compared with WT niche (**O**) is rescued in simultaneous loss of both the function of tak1 and Relish (**R**) whereas JNK activation was not observed in tak1 loss (**Q**) **S-V**. Increase in niche cell numbers observed upon loss of Relish from the niche (**T**) is rescued to control levels (**S**) in a simultaneous loss of both Relish and tak1 function from the niche (**V**) whereas no significant change in niche cell number was observed in tak1 loss. (**W**) Statistical analysis of the data in (**S-V**) (n=10, P-value = 6.9×10^−10^ for control versus Relish RNAi, P-value =1.8×10^−9^ for tak1^2^; Relish RNAi versus Relish RNAi; two-tailed unpaired Student’s t-test) The white dotted line mark whole of the lymph gland in all cases. In all panels age of the larvae is 96 hrs AEH. The nuclei are marked with DAPI (Blue). Individual dots represent biological replicates. Error Bar: Standard Deviation (S.D). Data are mean±s.d. *P<0.05, **P<0.01 and ***P<0.001.

Interestingly, activation of JNK alone (expression of Hep^act^) in the niche can recapitulate the phenotypes associated with Relish loss to a large extent, for example, hyperproliferative niche (visualized by Antp, ***Figure 5 figure supplement 1A-B’*** and ***C***), ectopic differentiation (visualized by Nimrod P1, ***Figure 5 figure supplement 1D-E’*** and ***F***) and upregulated cytoskeletal elements (visualized by Enabled, ***Figure 5 figure supplement 1G-H’’*** and ***I***). Moreover, downregulating wg function in the same genetic background restores the cell number within the niche. These results further validate the epistatic relation of JNK and Wg in context to the hematopoietic niche (***Figure 5 figure supplement 1J-M*** *and **N)***.

To further understand the relationship of Relish-JNK in the context of niche cell proliferation and functionality, a double knockdown of both JNK and Relish from the niche was analyzed. The concurrent loss of JNK and Relish rescues the increase in niche cell proliferation, which is seen upon Relish loss (***Figure 5D-G’*** and ***H***). Additionally, it restores the progenitor pool (***Figure 5 figure supplement 2A-D and E***) and rescues the precocious differentiation ***(Figure 5 figure supplement 2F-I and J)*** noted in the lymph gland upon Relish loss from the niche (***Figure 5E-E’*** and ***H***). Moreover, downregulating JNK in conjunction with Relish loss from the niche restores the abrogated filopodial extension (***Figure 5 I-L)***. The rescue in ectopic differentiation coupled with the resurrection of the filopodial extension suggests a re-establishment of the communication process between the niche and the progenitors. The quantitative analyses further reveal the restoration of filopodial length **(*Figure 5 M*)** and number ***(Figure 5N)*** in the above genotype.

To have a functional insight into this result, we checked the extracellular Hh level (Hh^Ext)^ in the same genetic background. We found that in the double knockdown of JNK and Relish, the level of Hh^Ext^ present in the progenitors is similar to that of the control (***Figure 5 figure supplement 1O-Q’*** and quantitated in ***Figure 5 figure supplement 1R-T***). Therefore, the downregulation of the elevated JNK in Relish loss restores niche cell number, as well as the proper communication between niche cells and progenitors, which is mandatory for the maintenance of the latter.

Collectively, these results indicate that Relish functions in the niche to repress JNK signaling during development. In the absence of this regulation, upregulated JNK causes cytoskeletal re-arrangements within the niche and disrupts Hh delivery to the progenitors. The morphogen trapped within the niche is unable to reach the progenitors, thereby affecting their maintenance.

### Relish inhibits JNK signaling by restricting *tak1* activity in the niche during development

It is essential to understand how in a developmental scenario, the repression on JNK by Relish is brought about. Several *in vitro* and *in vivo* studies in vertebrates have shown the inhibitory role of NF-κB signaling over JNK during various developmental and immune responses (Clark & Coopersmith, 2007; Nakano, 2004; Tang et al., 2001; Volk et al., 2014). In *Drosophila*, mammalian MAP3 kinase homolog TAK1 activates both the JNK and NF-κB pathways following immune stimulation (Boutros et al., 2002; Kaneko et al., 2006; Vidal, 2001). Interestingly during infection, Relish, once activated, leads to proteasomal degradation of TAK1, thereby limiting JNK signaling to prevent hyper-immune activation (Park, 2004). It is intriguing to speculate that a similar circuit is engaged in the niche to curtail JNK signaling during development. If this is the case, then the loss of *tak1* should restore the elevated TRE-GFP expression in a niche where Relish is downregulated. Indeed, upon genetic removal of one copy of *tak1* in conjunction with Relish loss from the niche, a drastic decrease in TRE-GFP expression is noted (***Figure 5O-R***). Further, on checking the niche cell proliferation, we found a significant reduction in cell number; analogous to what we observe when JNK and Relish activity is simultaneously downregulated from the niche (***Figure 5S-V*** and ***W***). It is interesting to note that there is a restoration in the progenitor number ***(Figure 5 figure supplement 2K-N and O)*** along with the rescue of the precocious differentiation ***(Figure 5 figure supplement 2P-S and T)*** observed upon Relish loss from the niche, which is comparable to the control state in the above genotype.

These results led us to infer that Relish restricts the activation of JNK signaling in the hematopoietic niche via *tak1* during development. The restraint on JNK activity is essential for proper communication between niche cells and progenitor cells, which is necessary for the maintenance of the latter.

### Ecdysone dependent activation of Relish in the niche is a developmental requirement

Cleavage, activation, and nuclear translocation of Relish during infection is brought about by binding of the cell wall component of gram-negative bacteria to membrane-bound receptor PGRP-LC (Kaneko et al., 2004; Leulier et al., 2003). We wondered whether the niche is employing a similar mechanism to regulate Relish activation during development by engaging the endogenous microbiota. To explore this possibility, we checked the status of the hematopoietic niche in the germ-free/axenic larvae (which were devoid of commensal microflora, ***Figure 6 figure supplement 1A-A’*** and ***B***). Compared to the control, we found no significant change in the niche cell number in an axenic condition (***Figure 6A-B’*** and ***D***). Additionally, JNK signaling (visualized by TRE-GFP) is not active in the hematopoietic niche of the axenic larva (***Figure 6 figure supplement 1C-C’***), neither the ectopic differentiation (visualized by Hemolectin, green) of the progenitors was evident (***Figure 6 figure supplement 1D-D’***). Further, we employed a deletion mutant allele of *PGRP-LB* (PGRP-LB delta). This gene codes for an amide that specifically degrades gram-negative bacterial peptidoglycan (PGN) (Paredes et al., 2011; Zaidman-Rémy et al., 2006). Even in this scenario, where systemic PGN level is known to be elevated there is no increase in the niche cell number (***Figure 6C-C’*** and ***D***). The above results demonstrate that during development, Relish expression and activation in the hematopoietic niche are independent of the commensal microflora.

**Figure 6:**
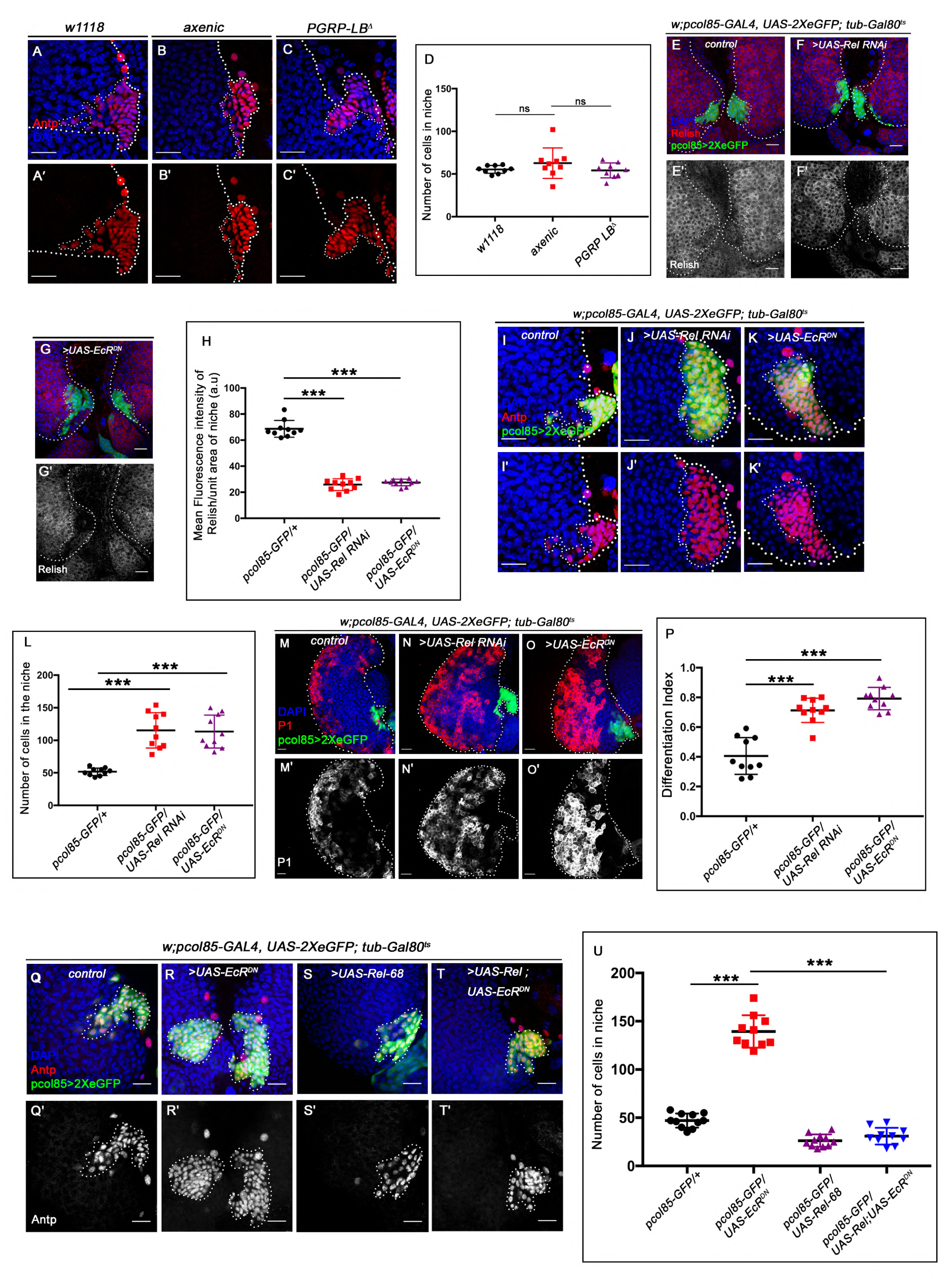
Ecdysone regulates Relish expression and functionality in the niche. The genotypes are mentioned in relevant panels. Scale bar: 20µm. **A-C’**. Niche number remains comparable to control (**A-A’**) both in axenic larval lymph gland (**B-B’**) and in PGRP-LB mutant where there is up regulation in systemic peptidoglycan levels (**C-C’**). (**D**) Statistical analysis of the data in **A-C’** (n=9; P-value = .262 for control versus germ free and .39209 for control versus PGRP-LB mutant; two-tailed unpaired Student’s t-test). **E-G’.** Compared to that of control (**E-E’**) Relish expression is significantly down regulated both in EcR loss (**G-G’**) as well as in Relish loss from the niche (**F-F’**). **H.** Statistical analysis of the data in **E-G’** (n=10, P-value =7.81×10^-12^ for control versus Relish loss and P-value = 3.76 x10^-10^ for control versus EcR^DN^; two-tailed unpaired Student’s t-test). **I-K’.** Similar to Relish loss from the niche (**J-J’**), EcR loss also results in increase in niche cell numbers (**K-K’**) compared to that of control (**I-I’**). **L**. Statistical analysis of the data in **I-K’ (**n=10, P-value = 6.6×10^-5^ for control versus EcR loss and P-value = 3.1×10^-5^ for control versus Relish loss; two-tailed unpaired Students t-test). **M-O’.** Compared to control (**M-M’**), both loss of Relish (**N-N’**) and EcR (**O-O’**) from the niche results in increase in differentiation. **P**. Statistical analysis of the data in **M-O’** (n=10, P-value = 4.2×10^-5^ for control versus Relish loss and P-value = 2.1×10^-6^ for control versus EcR loss; two-tailed unpaired Students t-test). **Q-T’.** Increase in niche cell numbers observed upon EcR loss from the niche (**R-R’**) is rescued to control levels (**Q-Q’**) when Relish was overexpressed in an EcR loss genetic background (**T-T’**). Overexpression Relish in the niche reduced the niche cell number compared to control (compare **T-T’** and **Q-Q’**). **U**. Statistical analysis of the data in **Q-T’** (n=10; P-value = 1.7×10^−9^ for control versus EcRDN, P-value = 7.8×10^−11^ for EcrDN versus UAS Rel; EcRDN, two-tailed unpaired Student’s t-test). The white dotted line mark whole of the lymph gland in all cases. In all panels age of the larvae is 96 hrs AEH. The nuclei are marked with DAPI (Blue). Individual dots represent biological replicates. Error Bar: Standard Deviation (S.D). Data are mean±s.d. *P<0.05, **P<0.01 and ***P<0.001.

Interestingly, activation of the IMD pathway components PGRP-LC and Relish is transcriptionally regulated by steroid hormone 20-Hydroxyecdysone signaling during infection (Rus et al., 2013). Moreover, a recent study also reveals that the activation of Relish, and IMD dependent genes is mediated via Ecdysone signaling in malphigian tubule during development (Verma & Tapadia, 2015). Strong expression of the Ecdysone receptor in the hematopoietic niche (***Figure 6 figure supplement 1E-E’’***) prompted us to check the possibility of ecdysone dependent regulation of Relish expression and activation in the niche. Upon expression of a dominant-negative allele of the receptor EcR in the niche, a drastic reduction in the amount of Relish protein is evident ***(Figure 6E-G’)***. Intensity analysis reveals a 3-fold decrease in Relish expression upon blocking ecdysone signaling compared to the control niches (***Figure 6H***). Since transcriptional regulation of Relish through ecdysone signaling has been previously reported (Rus et al., 2013), we decided to explore whether this holds in case of the hematopoietic niche. Fluorescent *in-situ* hybridization (FISH) analysis reveals the presence of *Rel* transcript in the lymph gland as well as the salivary gland of control third instar larvae (***Figure 6 figure supplement 2F-F’*** and ***H-H’***; ***Figure 6 figure supplement 2I-J*** :sense probe acts as negative control). Due to increase in differention, the number of *Rel* expressing progenitors are less compared to control (***Figure 6 figure supplement 2G-G’)*.**

To probe the status of Rel transcripts specifically in the niche, we performed whole-mount immunofluorescence (IF) along with fluorescent in-situ hybridization (FISH) on the third instar lymph gland. Drastic reduction of the *Rel* transcript is noticeable in the niche from where EcR expression was downregulated compared to the control ***(Figure 6 figure supplement 2K-L’’),*** implicating that *Rel* is transcriptionally regulated through ecdysone signaling.

This observation indicates that the phenotypes observed upon EcR loss from the niche should be analogous to Rel loss. Attenuation of ecdysone signaling indeed leads to a significant increase in niche cell proliferation compared to the control (***Figure 6I-K’*** and ***L***). Furthermore, to understand whether the functionality of the niche is also compromised in the above genotype, we checked the differentiation status. Similar to Relish loss, downregulation of ecdysone signaling from the niche results in precocious differentiation (***Figure 6M-O’*** and ***P***). Niche-specific overexpression of Rel in conjunction with EcR loss can restore the cell number of the niche (***Figure 6 Q-T’*** and ***U*** as well as its functionality (***Figure 6 figure supplement 2A-E***).

These results, therefore, collectively suggest that ecdysone signaling regulates the expression and activation of Relish in the hematopoietic niche during development (***Figure 6 figure supplement 1F***). These results also underscore the requirement of a hormonal signal in regulating Relish during developmental hematopoiesis.

### During infection Relish in the niche is downregulated to facilitate immune response

In *Drosophila*, ecdysone mediated immune potentiation has shown to have a greater impact on the development of immunity in embryos (Tan et al., 2014) as well as the survival of flies during infection (Flatt et al., 2008; Rus et al., 2013; Verma & Tapadia, 2015; Xiong et al., 2016). Interestingly, we found that there is a 4-fold decrease in Relish expression from the hematopoietic niche during bacterial infection compared to uninfected larvae (Compare ***Figure 7A-A’*** with ***C-C’*** and quantitated in ***Figure 7D***). To rule out the possible effect of injection on Rel expression, we compared the infected with sham control. There was a 2.6 fold decrease in the intensity of Rel expression within the niche of infected larvae compared to the sham control (Compare ***Figure 7B-B’*** with ***C-C’,*** quantitated in ***Figure 7D***). In contrast, upon infection, we could see the nuclear expression of Relish in the fat body cells as previously reported (***Figure 7E-G***) (Cha et al., 2003; Kim et al., 2006). Interestingly, niche-specific overexpression of N-terminal domain of Relish (*UAS-Rel68kD*), which is known to translocate to the nucleus and induce target gene expression (Stöven et al., 2000) is unable to sustain Relish expression post-infection (***Figure 7H-H’***) implicating the post-transcriptional regulation on Relish during infection. Relish activity is known to be modulated through proteasomal degradation in *Drosophila* and *Bombyx mori* (Khush et al., 2002; Ma et al., 2015).

**Figure 7:**
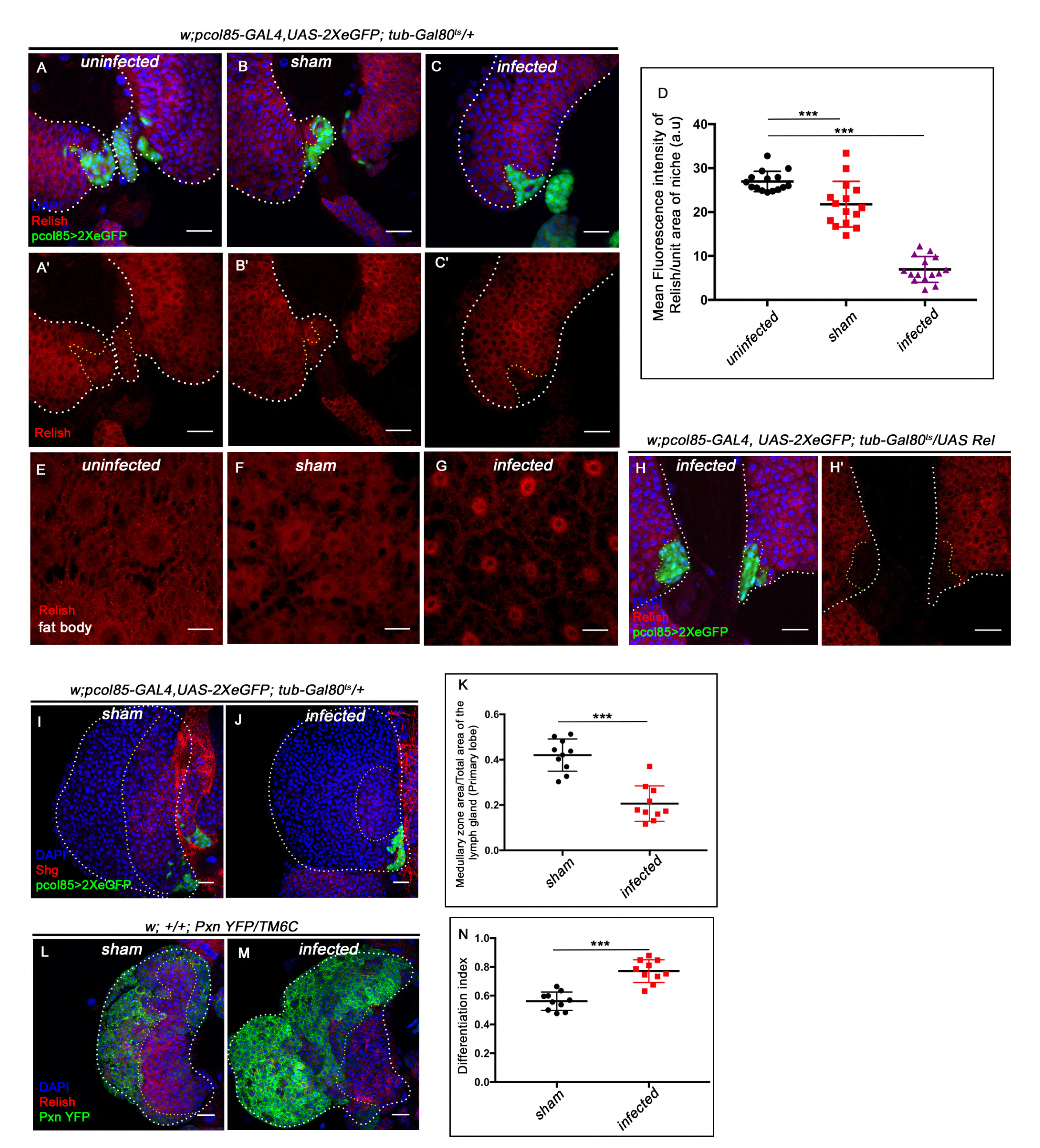
Niche specific expression and function of Relish is susceptible to pathophysiological state of the organism. The genotypes are mentioned in relevant panels. Scale bar: 20µm. **A-C’**. Compare to uninfected conditions (**A-A’**) and sham (**B-B’**), significant reduction in Relish expression was observed in the hematopoietic niche four hours post infection (**C-C’**). **D.** Statistical analysis of the data in **A-C’** (n=15; P= 6.62×10^−18^ for unpricked versus infected, P= 2.5×10^−7^ for sham versus infected, two-tailed unpaired Student’s t-test). **E-G.** Nuclear expression of Relish was observed in infected (**G**) fat body cells four hours post in contrast to uninfected (**E**) and sham (**F**) larval fat body. **H-H’.** Overexpressing Relish N-terminus (*UAS-Rel-68kD*) could not rescue loss of Relish expression post infection. **I-J.** Compared to sham (**I**), significant reduction in Shg positive progenitors (red) were observed in infected lymph glands (**J**). **K**. Statistical analysis of the data in **I-J** (n=10; P-value = 5.2×10^−6^ for sham versus infected, two-tailed unpaired Student’s t-test). **L-M.** Drastic increase in differentiation (visualised by Pxn-YFP, green) was observed in infected lymph glands **(L)** compared to sham **(M). N.** Statistical analysis of the data in **L-M** (n=10; P-value = 4.65×10^−6^ for sham versus infected, two-tailed unpaired Student’s t-test). The white dotted line mark whole of the lymph gland in all cases. In all panels age of the larvae is 96 hrs AEH. The nuclei are marked with DAPI (Blue). Individual dots represent biological replicates. Error Bar: Standard Deviation (S.D). Data are mean±s.d. *P<0.05, **P<0.01 and ***P<0.001.

More importantly, we also found that compared to control, four hours post-bacterial challenge, the progenitor pool declines (***Figure 7I-K***), accompanied by a concomitant precocious differentiation (***Figure 7L-N).*** These phenotypes show a remarkable similarity to the ones seen on the loss of Relish from the niche ***(Figure 1H-P***). As a response to systemic infection, upregulation of JNK is detected throughout the lymph gland, including the niche compared to sham control (***Figure 7 figure supplement 1A-B’).*** The short duration of systemic infection adopted in our study induced proliferation in the otherwise quiescent niche cells (***Figure 7 figure supplement 1C-E)***. Based on these studies, we speculate that Relish, in this case, might also undergo ubiquitin-mediated degradation (by Factor X, ***Figure 7F***) that overrides the developmental signal (***Figure 6 figure supplement 2F***) during infection.

These data collectively elucidate that a differential regulation on Relish is mandatory during infection to boost immune response.

## Discussion

Our study unravels how the hormonal control on Relish in the hematopoietic niche is essential for maintaining the hemocyte progenitors of the lymph gland during development. Hemocytes present in the lymph gland are not actively involved in immune surveillance under healthy conditions. Within this organ, the hemocytes proliferate to create a pool of progenitors and differentiated cells. However, with its content, this organ takes care of all post-larval hematopoiesis and therefore is not precociously engaged. Our study illustrates how the hematopoietic niche recruits neuroendocrine-immunity (Ecdysone-Relish) axis to maintain the progenitors of the lymph gland during larval development (***Figure 6 figure supplement 1F***).

The loss of Ecdysone/Relish, therefore, results in precocious maturation of the progenitors. The mechanism underlying the control of niche state and function by Relish involves repression of the Jun Kinase signaling. Interestingly, Relish during infection is known to inhibit JNK activation in response to gram-negative bacterial infection in *Drosophila* (Park, 2004). We found that this antagonistic relation of Relish and JNK, essential for innate immunity, is also relevant during development to facilitate the functioning of the hematopoietic niche. Our results suggest two independent events occur in the niche if JNK is activated (***Figure 6 figure supplement 1F***). Firstly, the activation of JNK leads to supernumerary niche cells due to an increase in Wingless expression. Secondly, the JNK pathway negatively regulates the actin-based cytoskeletal architecture essential for the release of Hh from the niche cells.

Though perceived as a pro-apoptotic signal, a large body of work has evidenced the role of the JNK pathway to induce proliferation in diverse developmental scenarios (Kaur et al., 2019; Ohsawa et al., 2012; Perez-Garijo et al., 2009; Pinal et al., 2019; Wu et al., 2010). The JNK pathway is also known for its ability to release proliferative signals that can stimulate the growth of the tissue (Pinal et al., 2019). For instance, during compensatory proliferation in the developing larval wing disc, JNK triggers wingless to stimulate the proliferation of the non-dead cells (Ryoo et al., 2004). Moreover, wingless signaling has been reported as a mitogenic signal for stem cells in diverse contexts (Deb et al., 2008; G. Lin et al., 2008; Song, 2003) and aberrant activation of this pathway contributes to various blood cell disorders and cancers (Grainger & Willert, 2018; Klaus & Birchmeier, 2008; Lento et al., 2013; Reya & Clevers, 2005). *Drosophila* hematopoietic niche is known to positively rely upon wingless (Wg) signaling for its proliferation during larval development. Down-regulation of the signaling by expressing a dominant-negative form of its receptor Frizzled results in a reduction in niche cell numbers (Sinenko et al., 2009). We believe, to prevent hyperproliferation of the niche cells, Relish is reining in Wingless by inhibiting JNK signaling during development.

Several studies have shown that actin-based cellular extensions or cytonemes (Bischoff et al., 2013; González-Méndez et al., 2019; Gradilla et al., 2014; Kornberg & Roy, 2014; Portela et al., 2019) play a crucial role in transporting Hh from the source to several cell diameter distances (Rojas-Ríos et al., 2012) thereby, contributing in the establishment of Hh gradient. Coincidently, *Drosophila* hematopoietic niche cells are also known to emanate cytoneme–like filopodial projections to the nearby progenitor cells (Mandal et al., 2007; Pennetier et al., 2012; T. Tokusumi et al., 2011). The current study is in sync with the understanding that these cellular extensions are required to maintain the undifferentiated cell population, possibly by facilitating the crosstalk between niche and hematopoietic progenitors (Krzemień et al., 2007; T. Tokusumi et al., 2011). Here, we show that upon Relish loss from the niche, filopodial formation gets impaired in a JNK dependent manner. Ectopic activation of JNK signaling leads to altered expression of cytoskeletal elements that disrupt the process of filopodial formation. Consequently, the morphogen Hh gets trapped within the niche cells thereby hampers the proper communication between the niche and the progenitor cells of the lymph gland (***Figure 8***). Previous studies have demonstrated that activation of Relish leads to the disruption of cytoskeletal architecture in S2 cells to bring about the necessary changes associated with cell shapes for the proper immune response (Foley & O’Farrell, 2004). However, the underlying mechanism of the modulation of cytoskeletal elements by Relish was not evident. Here we provide in vivo genetic evidence for the process by which Relish loss causes alteration of the cytoskeletal elements of the niche cells by ectopic JNK activation.

**Figure 8.**
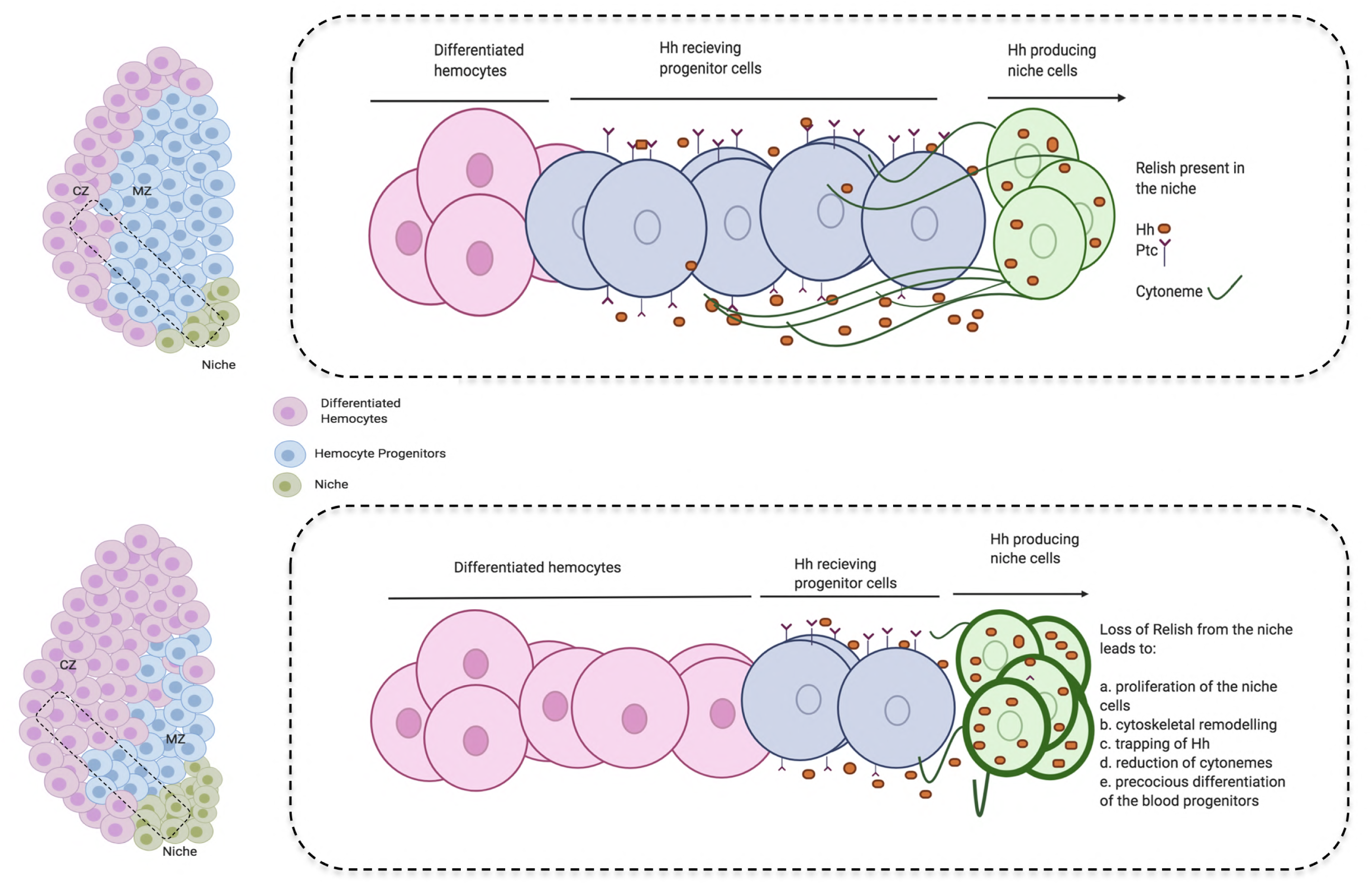
Developmental requirement of Relish in the niche for progenitor maintenance. Scheme describing how loss of Relish from the niche alters cytoskeletal elements of the cells. The change in cytoskeletal architecture affects cytoneme-like filopodial formation thereby trapping Hedgehog within the niche. The failure of Hh delivery in-turn interferes with progenitor maintenances and pushes them towards differentiation.

Another enthralling finding of our study is identifying 20-Hydroxyecdysone signaling as a regulator of Drosophila developmental hematopoiesis. The underlying reasons for this hormonal control on Relish seem to be intriguing. The need for this regulatory network during development may be related to the various microbial threats that are commonly confronted and dealt with by the circulating hemocytes of the larvae. While the circulating hemocytes cater to this need, the blood cells in the lymph gland proliferate and undergo maturation, creating a reservoir of hemocytes dedicated to deal with the post-larval requirements. Therefore, to safeguard the reserve population from responding to all of the common threats, the niche employs the Ecdysone-Relish axis to prevent the disruption in definitive hematopoiesis. However, during a high infection load, the lymph gland ruptures, suggesting a break in this circuit. This notion gets endorsed when the niche is analyzed post-infection. Quite intriguingly, we found that the Relish expression is downregulated from the niche, and the response of the lymph gland mimics the genetic loss of Relish from the niche. These observations confirm that the developmental pathway gets tweaked in the hematopoietic niche to combat high bacterial infection (***Figure 7 figure supplement 1F***).

During infection, the activation of Relish by ecdysone signaling in the fat body results in the production of antimicrobial peptides (Rus et al., 2013). We show that in contrast to this, upon infection, Relish needs to be downregulated in the niche in order to bolster the cellular immune response. This downregulation of Relish facilitates the release of a large pool of macrophages from the lymph gland to augment the circulating hemocytes to combat infection. The lymph gland hemocytes do not participate in the immune surveillance during development. However, during wasp infection, activation of the Toll/NF-κB signaling occurs in the niche to recruit lymph gland hemocytes to encapsulate wasp eggs (Louradour et al., 2017). We show that during bacterial infections Relish, another member of NF-κB pathway, is downregulated in the niche to disperse the lymph gland hemocytes into circulation. It is intriguing to see that the contrasting regulation of NF-κB components by the hematopoietic niche is essential for mounting an adequate immune response.

Interestingly, de novo production of neutrophils occurs in the bone marrow in response to systemic bacterial infection (Zhao & Baltimore, 2015). In mouse “emergency granulopoiesis” demands the activation of the TLR (Toll-like Receptors)/NF-κB pathway via TLR4 in the vascular niche (Boettcher et al., 2014). It will be important to investigate whether this differential regulation on NF-κB members is evident in vertebrate bone marrow niches during infection.

For an organism to combat an infection successfully, a quick shift of the ongoing hematopoiesis towards emergency mode is absolutely necessary. We show that the hematopoietic niche is the sensor that gauges the physiological state of the animal and diverts the basal hematopoiesis towards the emergency hematopoiesis.

In conclusion, the present work reveals an unexpected role of Relish in developmental hematopoiesis. Furthermore, it unravels the systemic regulation of the hematopoietic niche by the neuroendocrine system. Also, it sheds light on how during infection, this pathway gets suppressed to reinforce the cellular arm of the innate immune response.

## Materials and methods

### Key Resources Table

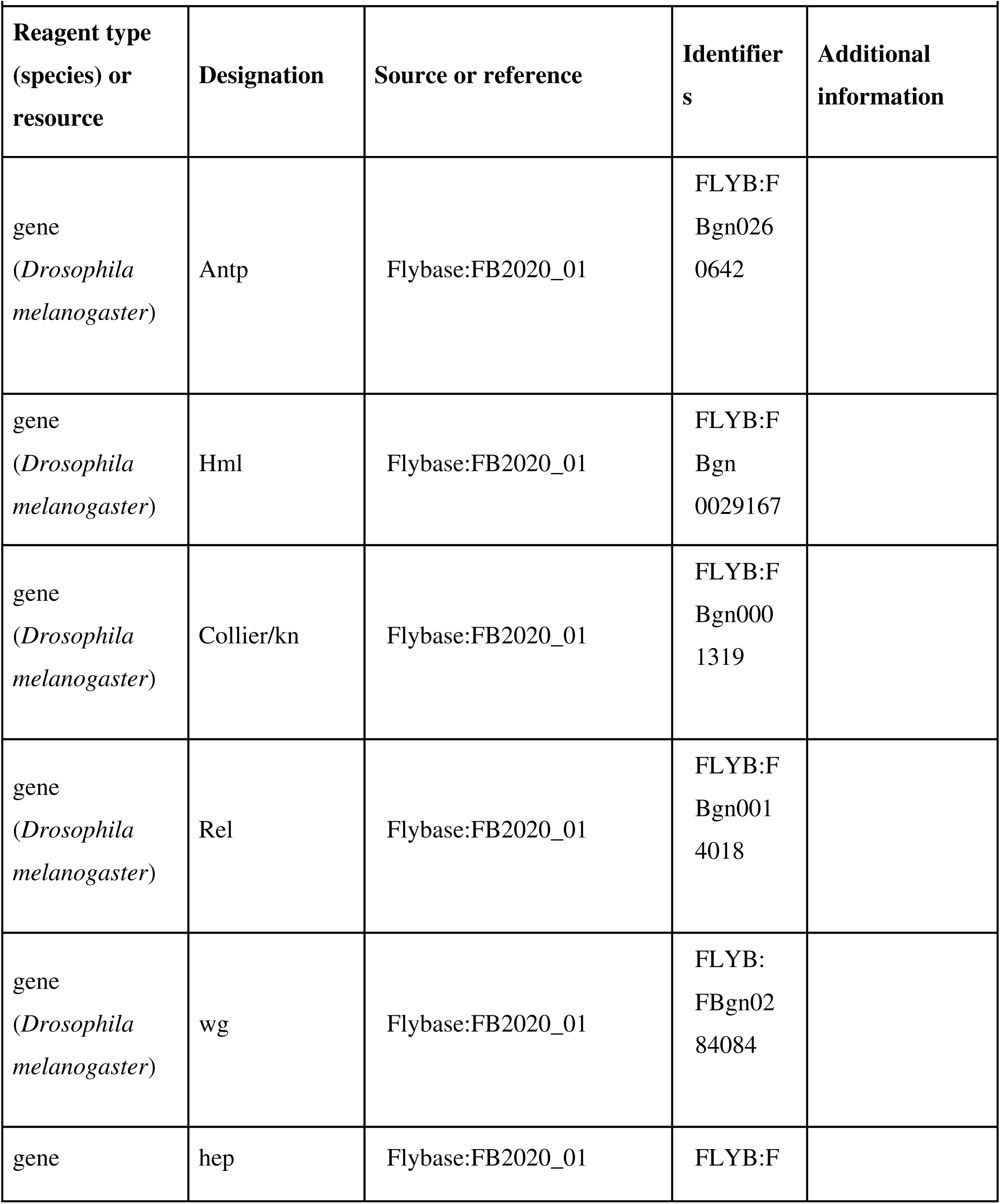
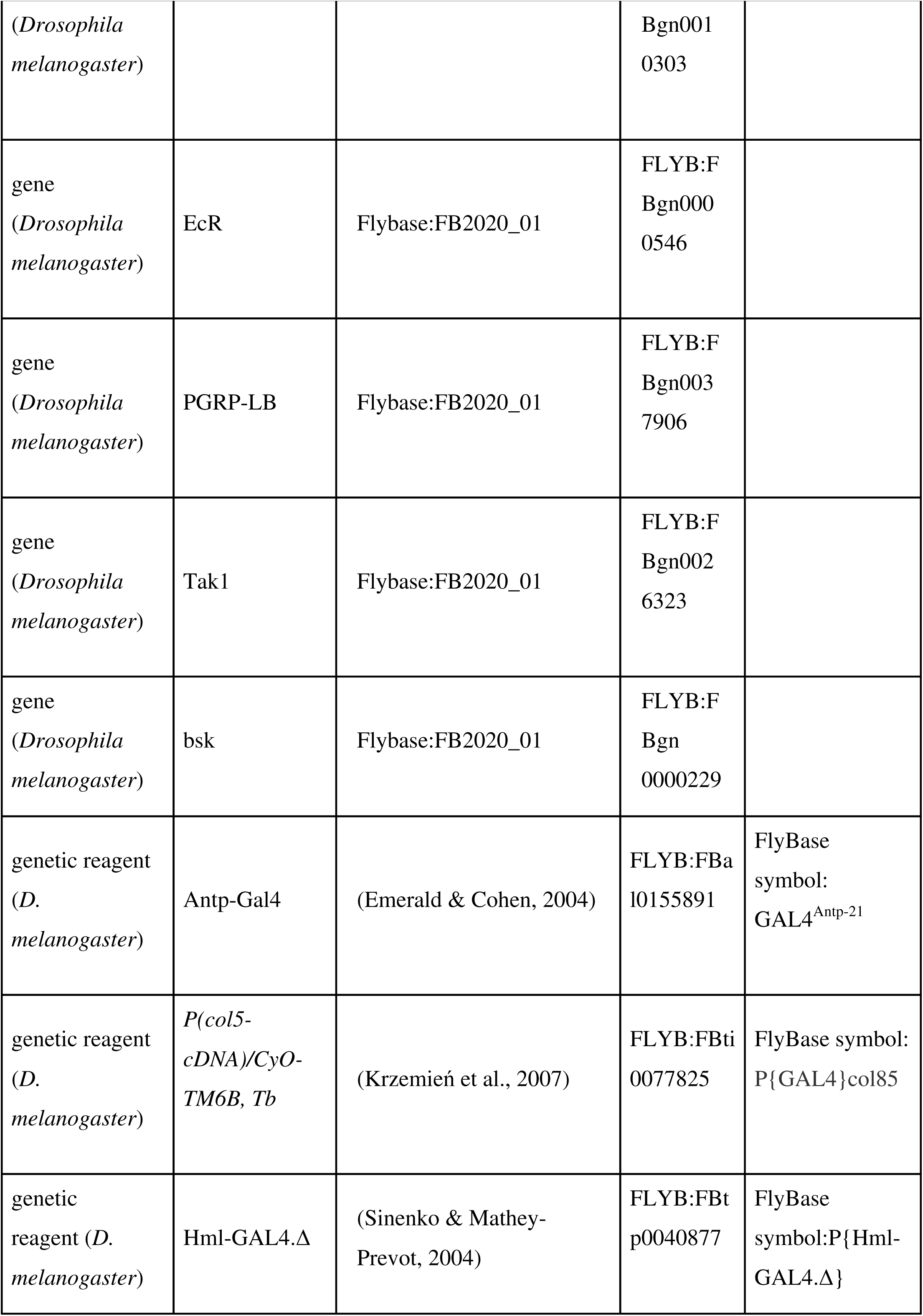
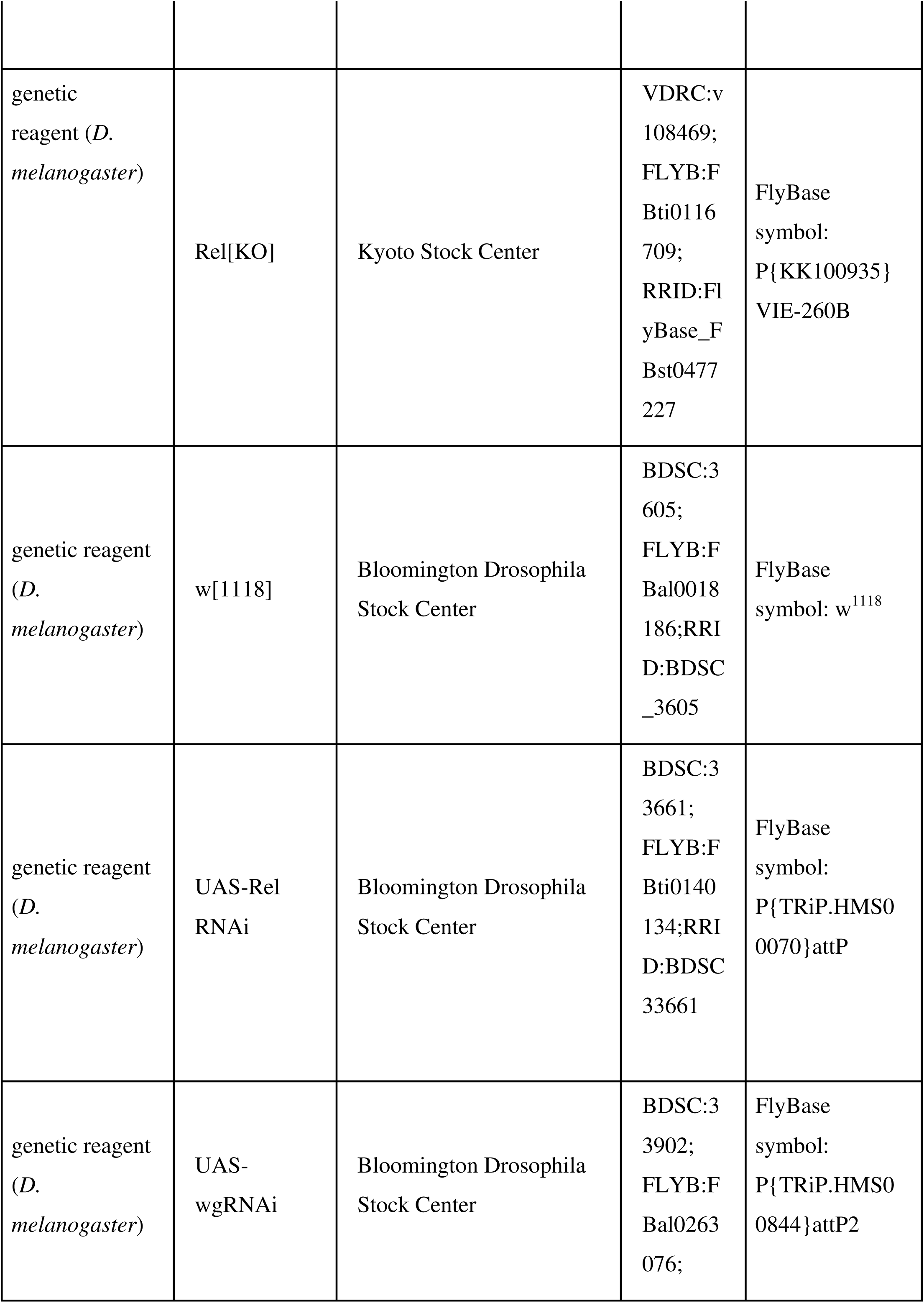
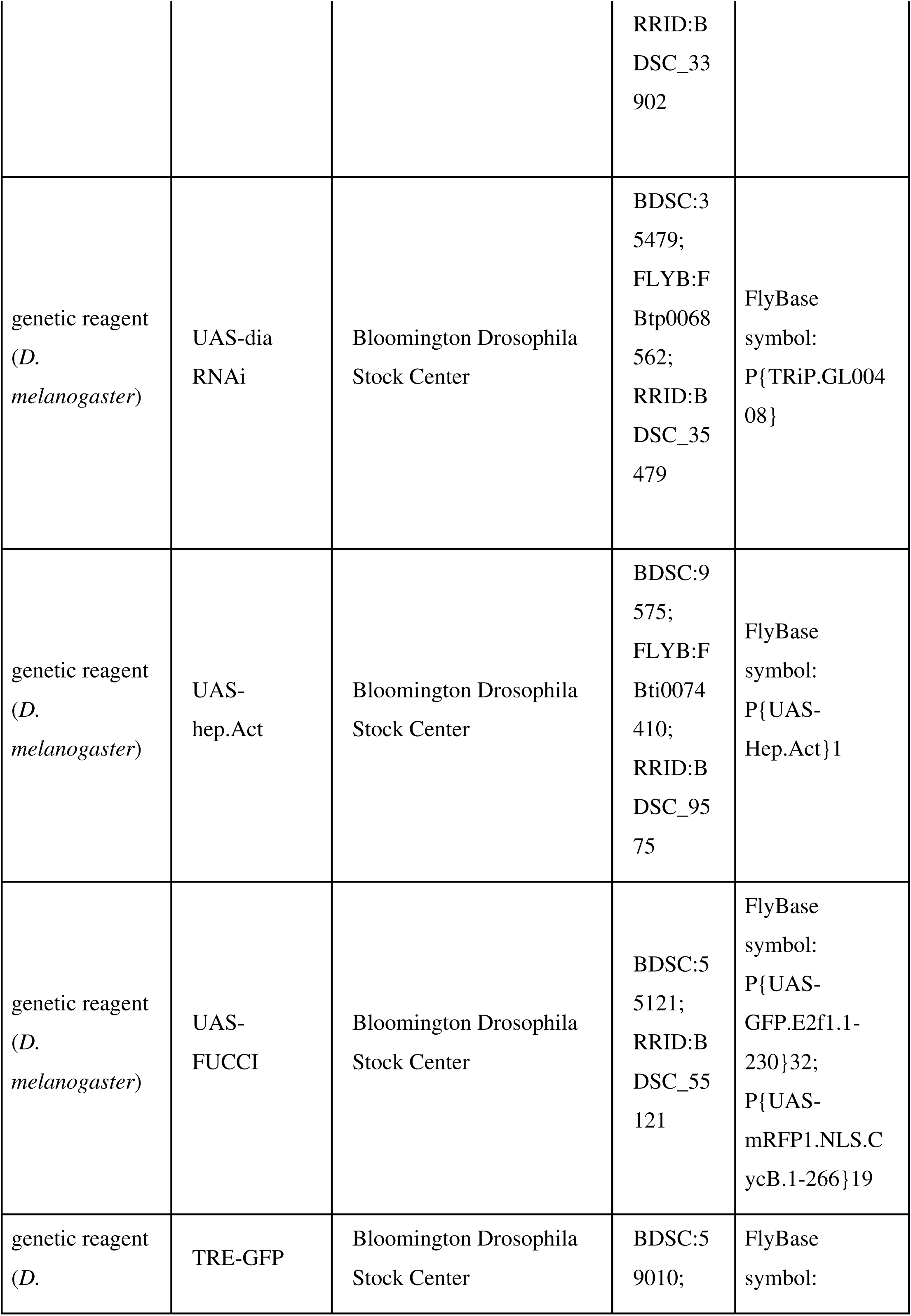
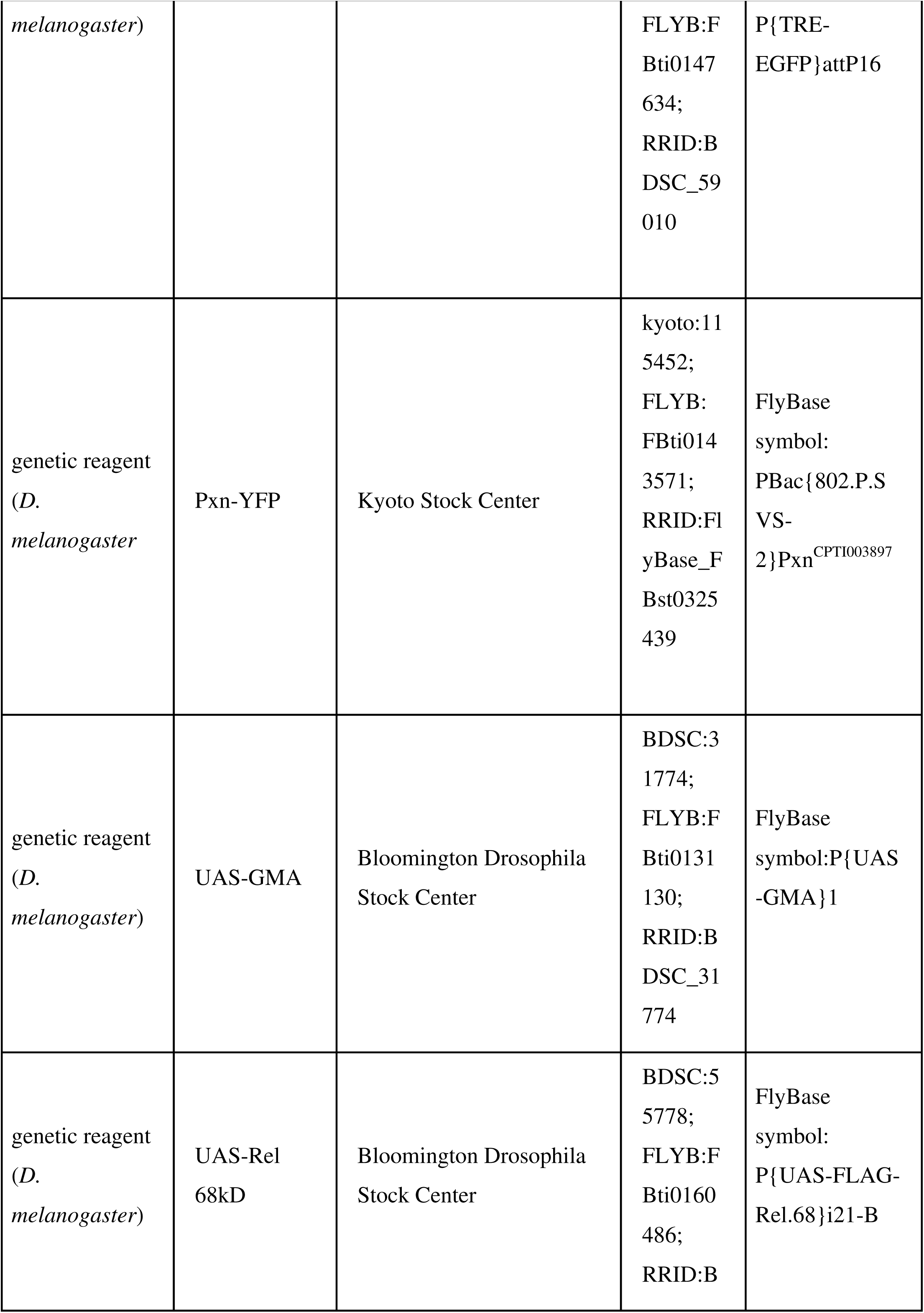
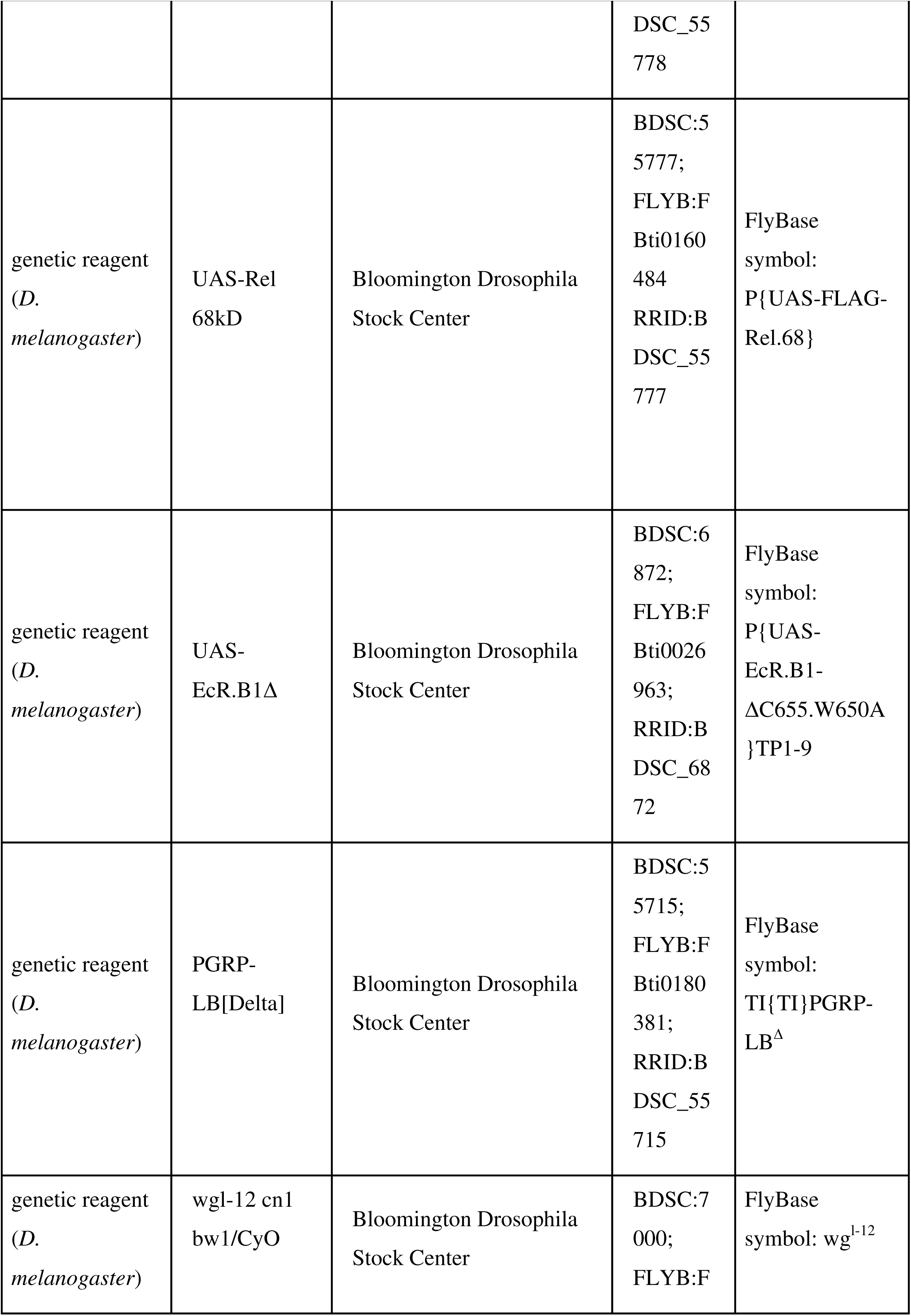
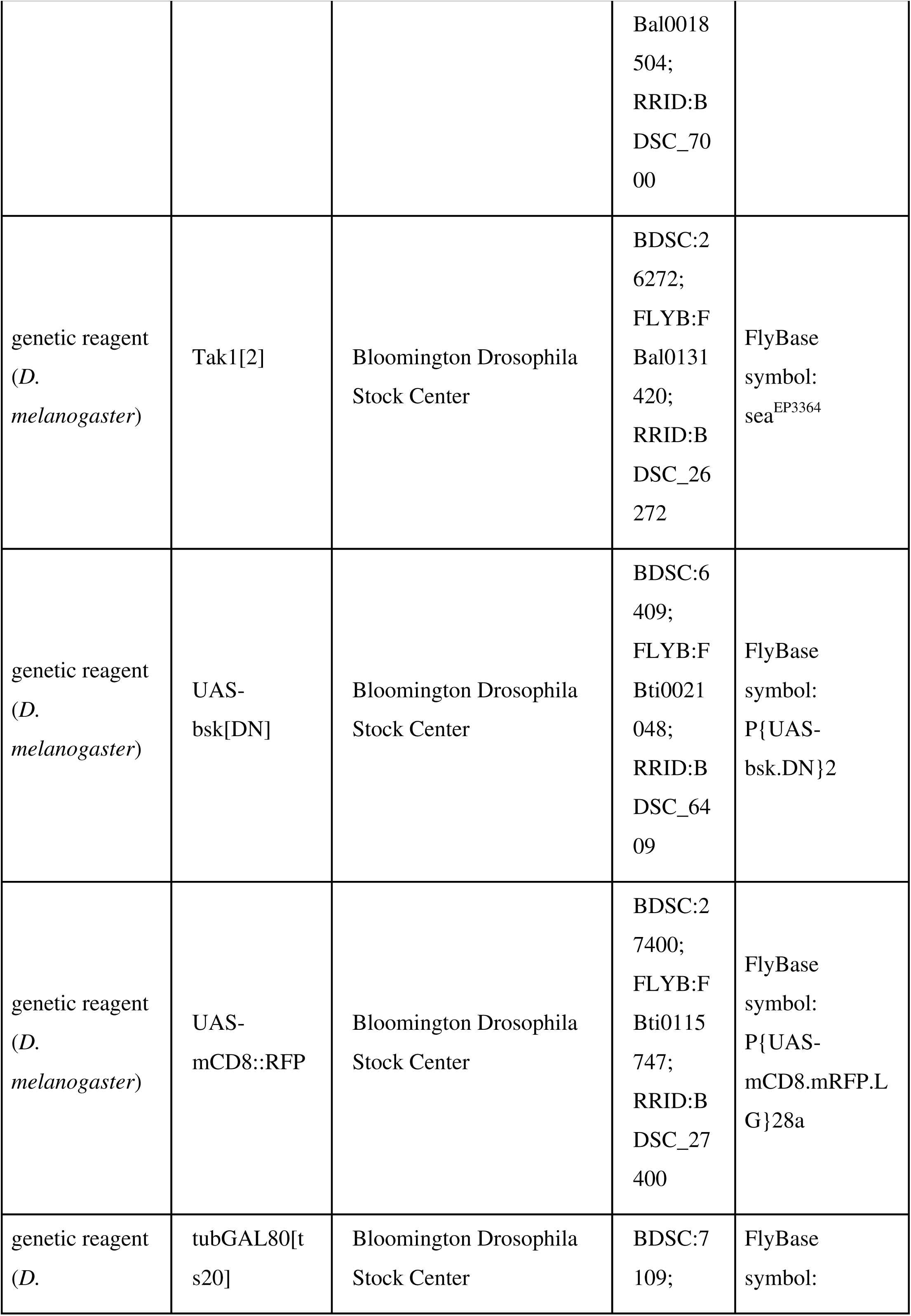
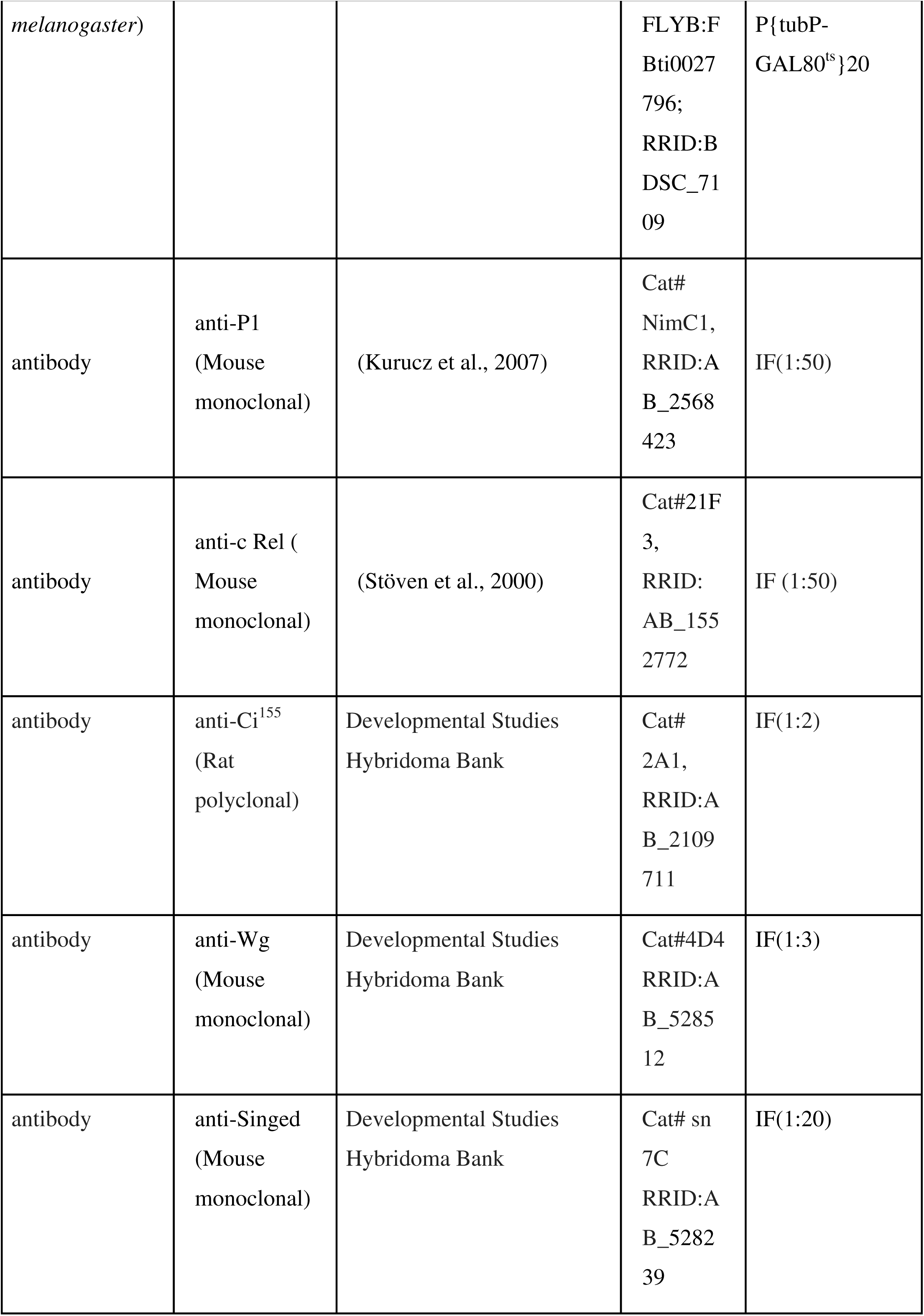
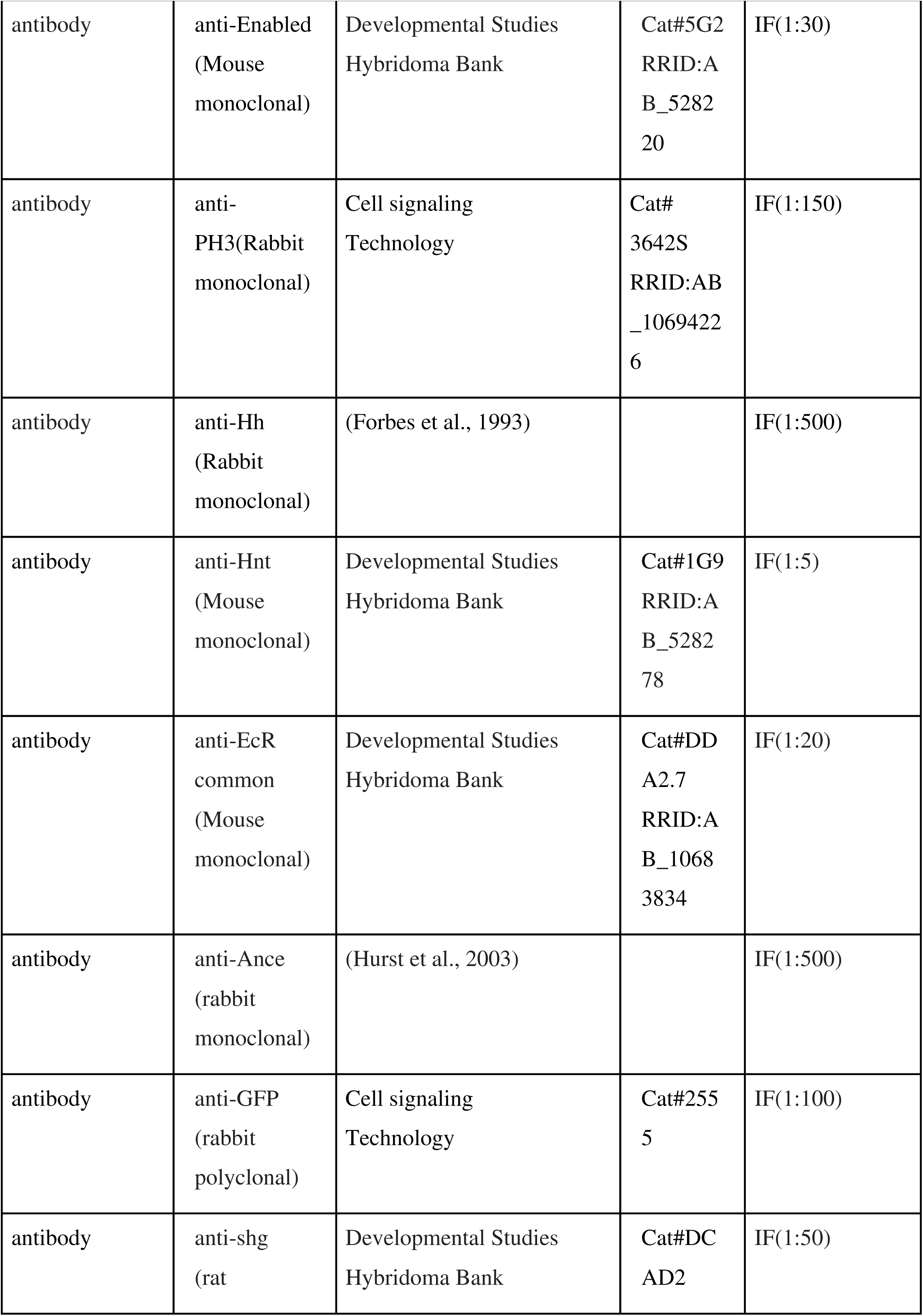
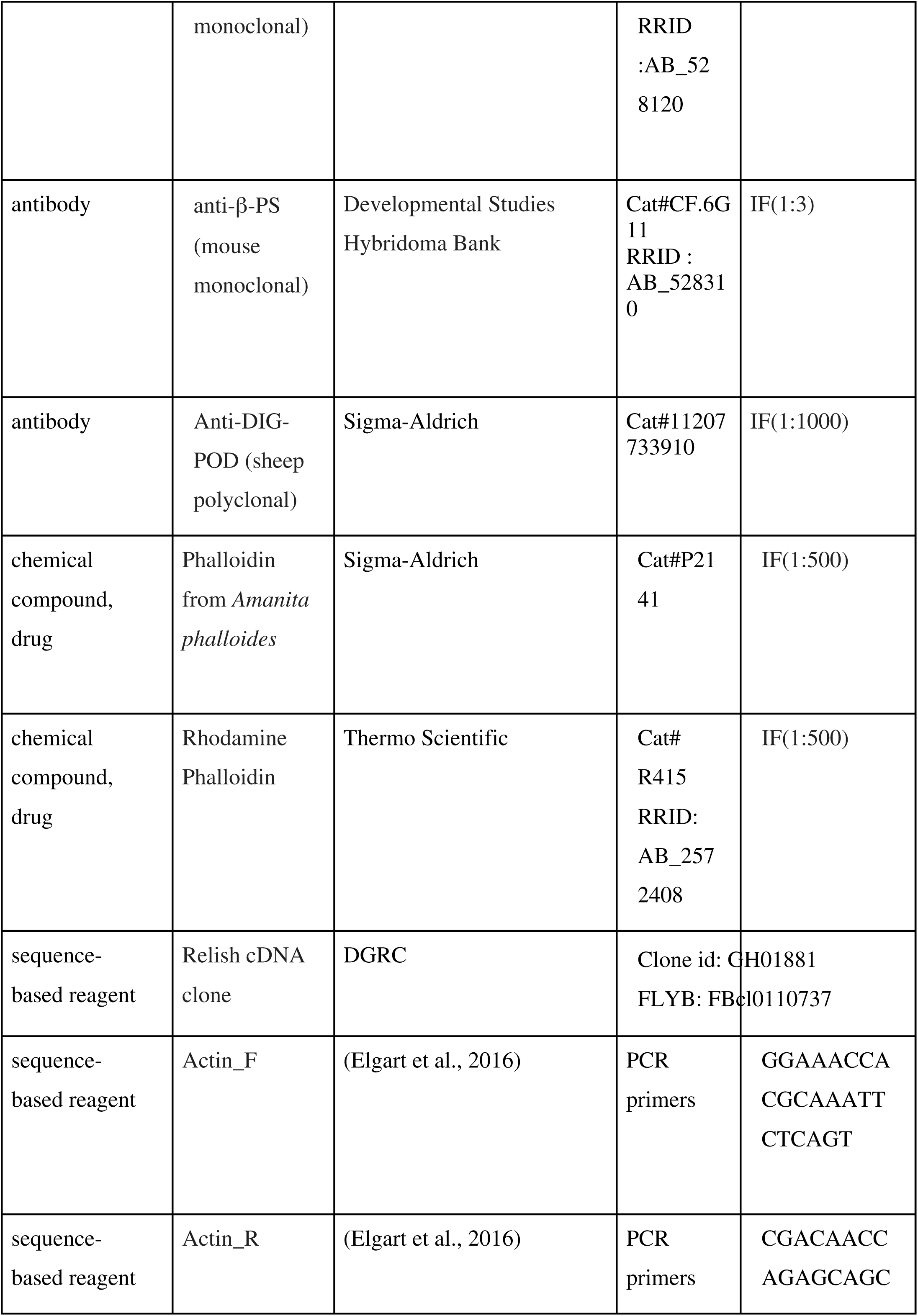
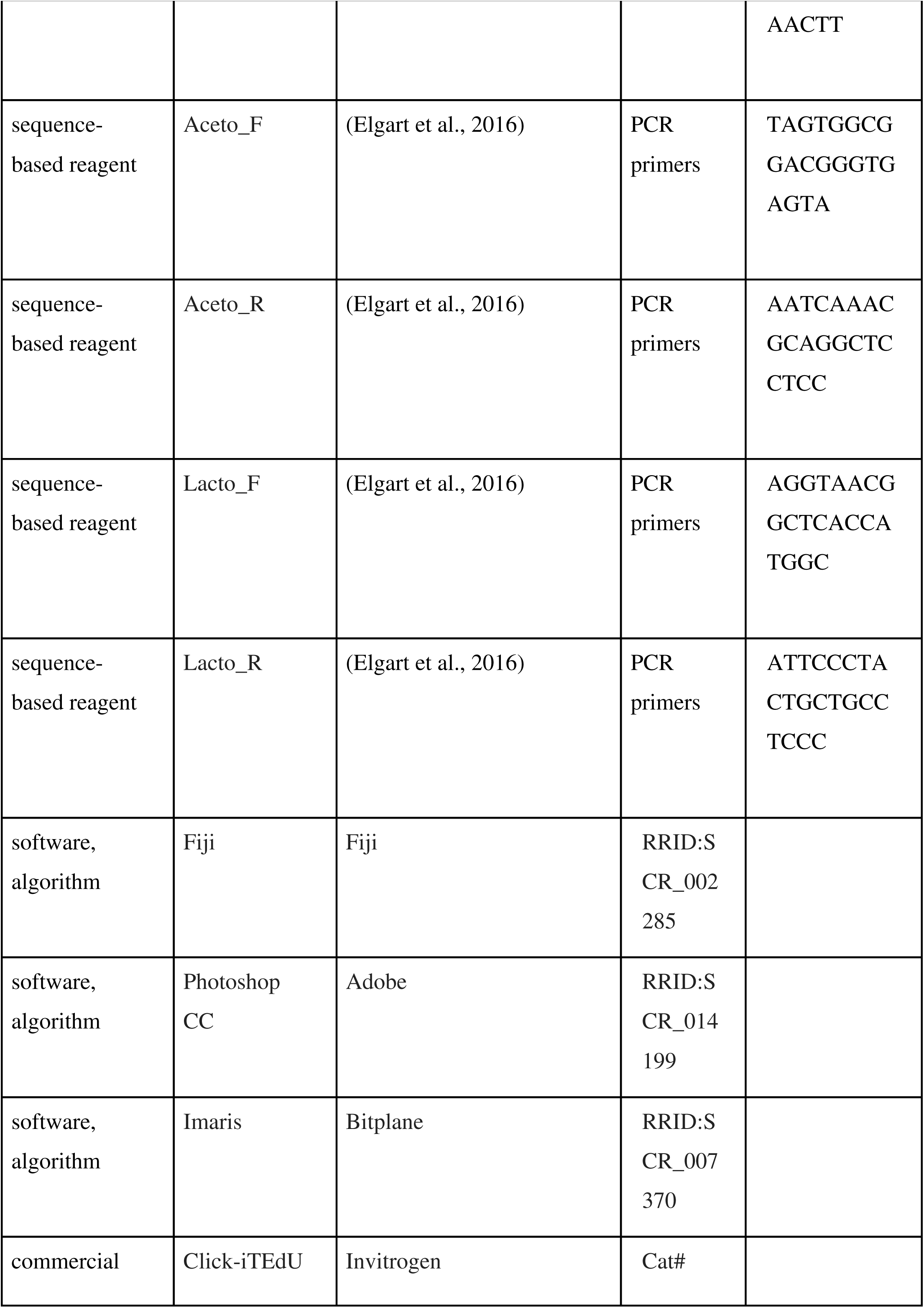
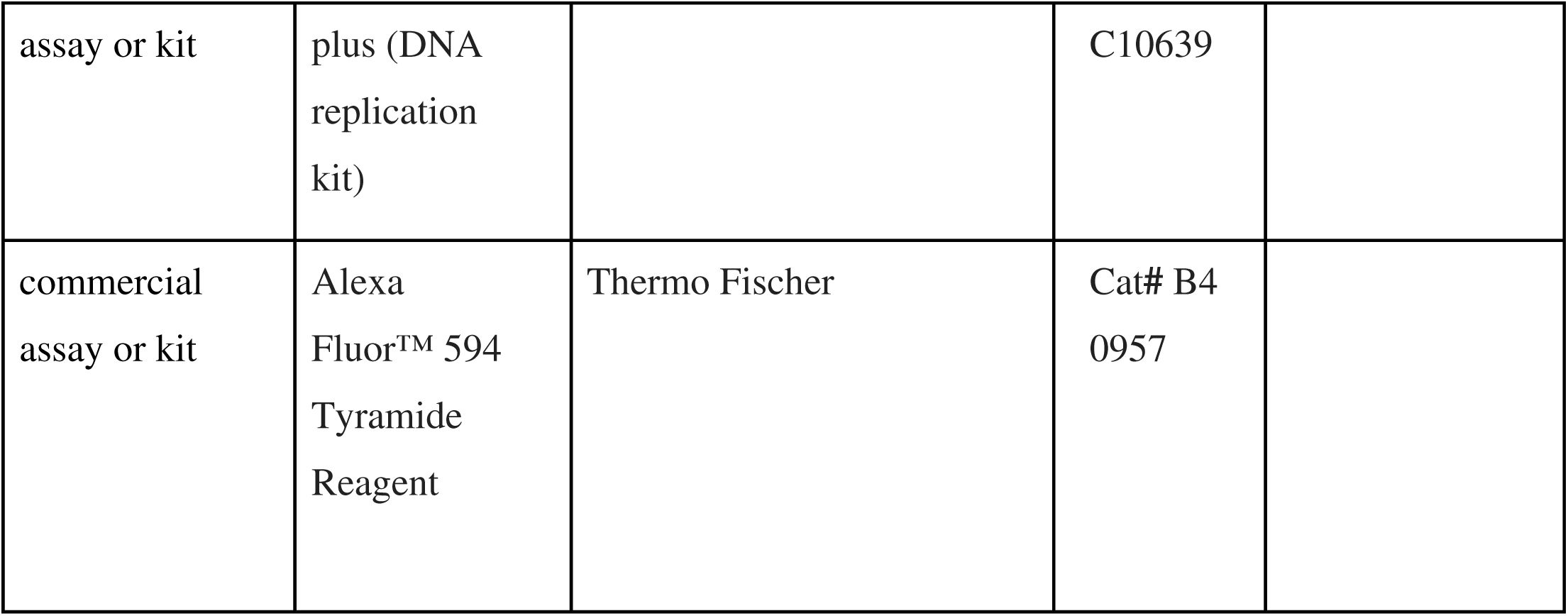

**Following genotypes were recombined for the current study:-**

1. Antp-Gal4.UAS-mCD8-RFP/Tb
2. TRE-GFP/TRE-GFP; Antp-Gal4.UAS-mCD8-RFP/Tb
3. UAS bsk^DN^/UAS bsk^DN^; +/+; UAS Relish RNAi/UAS Relish RNAi
4. UAS-GMA/UAS-GMA; tubgal80^ts^/ tubgal80^ts^; Antp-Gal4 /Tb
5. w;pcol85-Gal4, UAS-2XeGFP; tub-Gal 80^ts^
6. UAS Relish RNAi^KK^/UAS Relish RNAi^KK^; UAS Wg RNAi/ UAS Wg RNAi
7. tubgal80^ts^/ tubgal80^ts^; Antp-Gal4.UAS-2XEGFP/TM2
8. UAS Relish /UAS Relish; UAS EcR DN/ UAS ECR DN.

All stocks were maintained at 25°C on standard media. For *GAL80^ts^* experiments, crosses were initially maintained at 18%°C (permissive temperature) for 2 days AEL to surpass the embryonic development, and then shifted to 29%°C till dissection.

For time series experiments, synchronization of larvae was done. Flies were allowed to lay eggs for about 4 hours. Newly hatched larvae within one-hour intervals were collected and transferred onto food plates and kept at 29°C till dissection.

### Immunohistochemistry

Immunostaining and dissection (unless said otherwise) were performed using protocols described in (Jung, 2005; Mandal et al., 2007; Mondal et al., 2011) using primary antibodies: mouse anti-c-Rel (1:50, a gift from N.Silverman (Stöven et al., 2000), mouse anti Relish (1:50, 21F3, DSHB), mouse anti-Antp (1:10, 8C11, DSHB), mouse anti-Wg (1:3, 4D4, DSHB), mouse anti-P1 (1:40, a gift from I. Ando, rabbit anti-Ance (1:500, a gift from A. D. Shirras), rat anti-Ci (1:5, 2A1, DSHB), mouse anti-singed (1:20, Sn7C, DSHB), mouse anti-enabled (1:30, 5G2, DSHB), rabbit anti-PH3 (1:150, Cell signaling), rabbit anti-Hh (1:500, a gift from P. Ingham), mouse anti-Hindsight (1:5, 1G9, DSHB), mouse anti-EcR common (1:20, DDA2.7, DSHB), mouse anti-β-PS (1:3, CF.6G11, DSHB), rabbit-anti-GFP(1:100, 2555, Cell signalling), rat anti-shg (1:50, DCAD2, DSHB). Secondary antibodies used in this study are as follows: mouse Cy3, mouse FITC, mouse Dylight 649, rabbit Cy3, (1:500) rabbit FITC, (1:200), Jackson Immuno-research Laboratories.

Tissues were mounted in Vectashield (Vector Laboratories) then followed by Confocal Microscopy (LSM, 780, FV10i, LSM 900).

### EdU Incorporation assay

Click-iT EdU (5-ethynyl-2’-deoxyuridine, a thymidine analog) kit from Life Technologies was used to perform DNA replication assay (Milton et al., 2014). Larval tissue was quickly pulled out in 1X PBS on ice (dissection time not more than 25 min and fat body and salivary gland needs to be cleared from the tissue of interest). Incubation of the dissected tissue was done in EdU solution, Component A (1:1000) in 1X PBS on shaker at room temperature for 30-35 minutes followed by fixation in 4% paraformaldehyde (prepared in 1XPBS). Post fixation tissues were washed with 0.3% PBS-Triton four times at ten minutes interval followed by 30-35 minutes of blocking in 10% NGS in 0.3% PBS-Triton. EdU staining solution as per manufacturer’s instruction (for 50 µ l staining solution, 43 µl 1x EdU buffer, 2 µl CuSO_4_ solution, 5 µl 1x EdU buffer additive, 0.12 µl Alexa solution) was used to stain the sample for 30 min at room temperature. Two quick washes with 0.3% PBS-Triton was followed by a quick wash in 1xPBS. If no further antibody staining was required, nuclear staining by DAPI was done in 1XPBS and then mounted in Vectashield.

### Extracellular Hh staining and quantitation

For extracellular Hh staining, a *detergent*-free *staining* protocol was used. Lymph glands were dissected in ice-cold Schneider’s media (Gibco 21720024, rinsed with cold PBS twice, and fixed with 4% formaldehyde overnight at 4°C (Sharma et al., 2019). Subsequent processing of the samples was the same as mentioned above in the Immunohistochemistry section, except that no detergent was used.

Protocol described by (Ayers et al., 2010) was used as a reference to perform quantitation. A rectangle (500 x 150 pixels) was drawn, spanning from the niche to the cortical zone diagonally with the medullary zone in the middle, as shown in Figure 4F. An extracellular Hedgehog profile was made using the “Plot Profile” tool of ImageJ. The Plot profile tool displays a “column average plot”, wherein the x-axis represents the horizontal distance through the selection and the y-axis the vertically averaged pixel intensity, which in this analysis is formed by extracellular Hedgehog staining.

### Filopodial detection and quantitation

*UAS-GMA* was used to label the filopodia using a niche-specific driver, *Antp-GAL4*. Lymph glands of the desired genotype were dissected in Schneider’s media (Gibco 21720024) and incubated in a solution containing Schneider’s media supplemented with 1% Phalloidin from *Amanita phalloides* (P2141 SIGMA) for 15 minutes in order to stabilize the filopodia. These tissues were then mounted and imaged directly under the confocal microscope.

Filopodial quantitation was done using ImageJ. The number of filopodia emanating from the niche in all the Z-stacks was counted manually per sample. The average number of filopodia emanating per sample was plotted using GraphPad Prism for different biological replicates. For filopodial lengths, the “Freehand line” tool was used to mark the entire filopodial lengths, and the “Measure” tool was employed to get values in µM. Filopodial lengths in all samples were then plotted collectively as individual points in GraphPad Prism.

### Phalloidin Staining

Lymph glands dissected were fixed and incubated in rhodamine-phalloidin (1:100 in PBS) (Molecular Probes) for 1hr. The samples were then washed thrice for 10 min in PBS followed by mounting in DAPI Vectashield before imaging.

### Quantification of Intensity Analysis of Phalloidin

Membranous intensity of Phalloidin was measured using line function in Image J/Fiji. Mean intensity was taken in a similar manner as mentioned in (Jiwon Shim et al., 2012) P values of <0.05; <0.005 and <0.0005, mentioned as *, **, *** respectively are considered as statistically significant.

### Imaging and Statistical Analyses

All images were captured as Z sections in Zeiss LSM 780 confocal microscope and Olympus Fluoview FV10i (Panel 7). Same settings were used for each set of experiments. All the experiments were repeated at least thrice to ensure reproducibility. Mostly, 10 lymph glands were analysed per genotype for quantification analysis. Data expressed as mean+/-Standard Deviation of values from three sets of independent experiments. At least ten images of the lymph gland /niche were analysed per genotype, and statistical analyses performed employed two-tailed Student’s t-test. P-values of <0.05; <0.01 and <0.001, mentioned as *, **, *** respectively are considered as statistically significant. All quantitative analysis was plotted using GraphPad.

### Quantitative analysis of cell types in Lymph Gland: PSC Cell Counting

Antp positive cells were counted using the spot function in imaris software (Sharma et al., 2019). Data from three independent experiments are plotted in GraphPad prism as mean+/-standard deviation of the values. All statistical analyses performed employing two-tailed Student’s t-test. http://www.bitplane.com/download/manuals/QuickStartTutorials5_7_0.pdf).

### Quantification of Intensity Analysis

Intensity analysis of Hh, TRE-GFP, Wg, Singed, Enabled and Relish in different genotypes was done using protocol mentioned in from (J. Shim et al., 2013) http://sciencetechblog.files.wordpress.com/2011/05/measuring-cell-fluorescence-using-imagej.pdf. For each genotype, in about ten biological samples, at least five ROIs were quantified. Data is expressed as mean+/-Standard Deviation of values and are plotted in GraphPad prism. All statistical analyses performed employing a two-tailed Student’s t-test.

### Differentiation index calculation

To calculate the differentiation index, middle most stacks from confocal *Z* sections were merged into a single stack for each lymph gland lobe using ImageJ/Fiji (NIH) software as described earlier (J. Shim et al., 2013). P1 positive area was marked by using Free hand tool. The size was measured using the Measure tool (Analyse–Measure).In similar way DAPI area was also measured. The differentiation index was estimated by dividing the size of the P1 positive area by the total size of the lobe (DAPI area). For each genotype, mostly 10 lymph gland lobes were used and Statistical analysis was performed using two tailed Student’s t test.

### *Fucci* cell cycle analysis

*UAS-GFP-E2f1_1-230_ UAS-mRFP1NLS-CycB_1-266_* (Zielke & Edgar, 2015) *f*ly line depends on GFP and RFP tagged degrons from E2F1 and Cyclin B proteins. Both E2F1 and Cyclin B gets degraded by APC/C and CRL4^cdt2^ ubiquitin E3 ligases once they enter S and G2-M phase of cell cycle respectively. Due to accumulation of *GFP-E2f1_1-230_*G1 phase will show green fluorescence and due to accumulation of *mRFP1NLS-CycB_1-266_S phase will show red fluorescence. Since bothGFP-E2f1_1-230_ and mRFP1NLS-CycB_1-266_are present in G2 and M phase, the cells will show yellow fluorescence.UAS-GFP-E2f1_1-230_ UAS-mRFP1NLS-CycB_1-266_* fly stock was recombined with *Antp-Gal4* and was crossed to *UAS-Relish RNAi* and *w^1118^* to ascertain the cell cycle status niche cells

All flies were kept at 25°C and Larvae were dissected 96 hr AEH.

### Generation of axenic batches

Germ free batches were generated following the ethanol based dechorination method provided in (Elgart et al., 2016). According to this method, embryos were collected, washed using autoclaved distilled water to get rid of residual food particles. Embryos were further dechorinated for 2-3 minutes in 4% sodium hypochlorite solution. Once this is done embryos were washed with autoclaved distilled water and were transferred to the sterile hood. Further manipulations were done inside the hood in order to avoid cross-contamination. Embryos were further washed twice with sterile water and were transferred into standard cornmeal food supplemented with tetracycline (50 microgram/ml).

### Bacterial plating experiment

For plating experiments, 3-5 late third instar larvae were washed in 70% ethanol twice for 2 min. Further, the larvae was washed using sterile H_2_O twice for 2 min. After this surface sterilization, the larvae were transferred into LB media and were crushed thoroughly using a pestle. Once crushed the homogenates were spread on LB agar media and was incubated for 3-4 days at 25°C.

### Measuring of Bacterial content by qPCR

To measure bacterial composition in the gut, 12–15 3rd instar larval guts were dissected and pooled and DNA was isolated manually using the protocol provided by VDRC (https://stockcenter.vdrc.at/images/downloads/GoodQualityGenomicDNA.pdf) followed by PCR analysis using species-specific primers. *Drosophila* actin was used as a control.

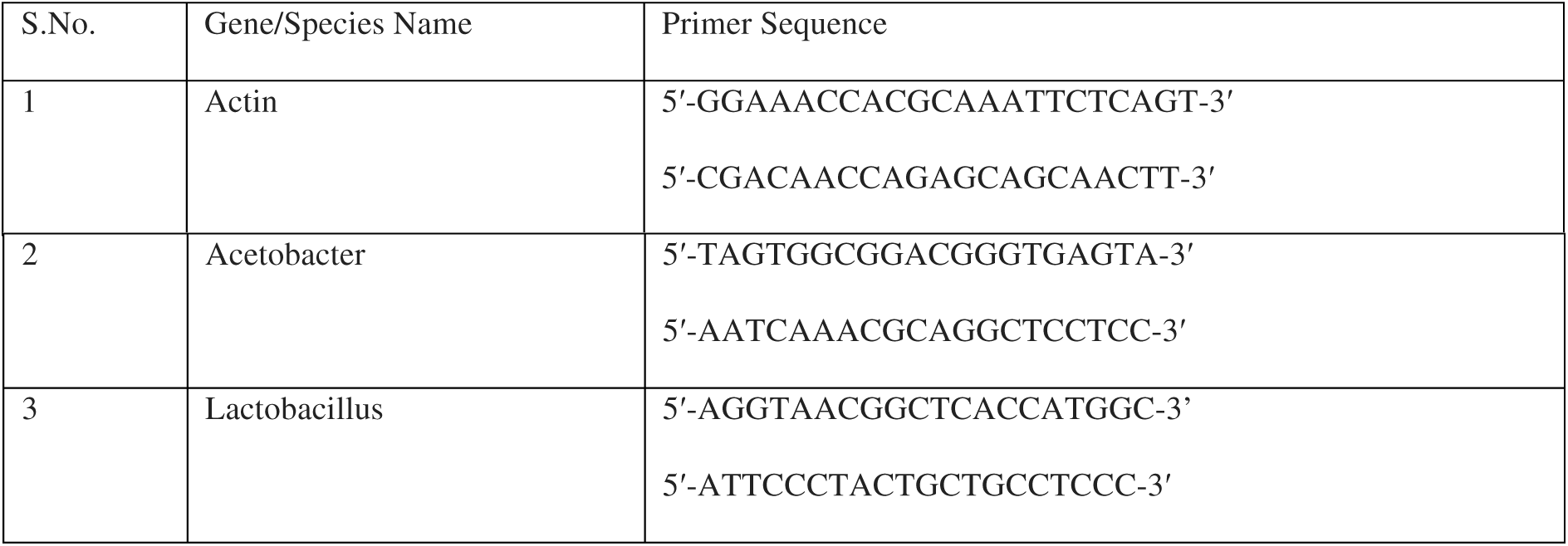

### Infection experiments

The following bacterial strains were used for infection: *E.coli* (OD_600_:100). For larval infection, bacterial cultures were concentrated by centrifugation; the pellet formed was resuspended in phosphate-buffered saline (PBS) to appropriate OD value. Synchronized third instar larval batches were used for all analyses. Third instar larvae were washed three times with sterile ddH_2_O and pricked using a fine insect pin dipped in bacterial suspension at the postero-lateral part. Mock injections were done using PBS dipped pins. Complete penetration was confirmed while dissection by looking at the melanization spots at the larval epithelial surface. Once infected, larval batched were transferred to food plates and incubated at 25^0^Celsius till dissection. All observations were made 4 hours post-infection.

### IF-Fluorescence In Situ hybridisation

The protocol we followed was modified from (Toledano et al., 2012)

#### a. Probe preparation

Rel clone was procured from DGRC. Following plasmid linearization and restriction digestion using EcoRV and Xho1, the DNA fragments were loaded in agarose gel for electrophoresis. Further, the desired DNA fragments were purified using PCI (Phenol: chloroform: isoamyl alcohol) based gel purification and DIG-labelled RNA anti-sense and sense probe was prepared using Sp6 and T7 polymerase enzyme, respectively. Following DNase treatment, the probes were precipitated using Licl_2_ and ethanol. The RNA pellet was dried resuspended in RNase-free dH_2_0, and stored at −80^0^C till further use.

#### b. Dual Labelling of mRNA and protein in the hematopoietic niche

For IF-FISH, we followed the Part B of the Tissue preparation and fixation section of (Toledano et al., 2012). Followed by quick dissection, the larval tissues (make sure of having minimum fat body cells since it can hinder the fixation and hybridization) were fixed for 30 minutes in 4% formaldehyde prepared in RNase free PBS, further washed in PBTH ((PBS containing 0.1% Tween 20 and 250 µg/ml yeast tRNA) for thrice, 10 minutes each. Samples were blocked using 5% BSA prepared in PBTHR (PBTH containing 0.2 U ml ^−1^ RNase inhibitor and 1 mM DTT). Further, tissues were incubated in rabbit anti-GFP (1:100, prepared in PBTHR) for 18 hours at 4^0^C. Tissues were washed using PBTH three times 10 minutes each, followed by blocking for 30 minutes using 5% BSA prepared in PBTHR. The tissues were then incubated in fluorescent-labeled secondary antibody (rabbit-FITC 1:100) for 4 hours at room temperature in a shaker. Following this, three washes of PBTH, 10 minutes each, tissues were fixed using 10% formaldehyde for 30 minutes. Post fixation, tissues were washed thrice, 5 minutes each and rinsed with 0.5 ml of prewarmed Hybridization buffer (HB) for 10 min in a 65 °C in a hybridisation chamber. Tissued were then blocked with PHB (HB mixed with tRNA (10mg/ml)) for 1 h in 65 °C. Following blocking, tissues were transferred to preheated RNA probe prepared in PHB (2 µg/ml) and incubated at 65 °C for 18 hours. Post hybridization, stringent washes were given using 0.1% PBT: HB mix as mentioned in Toledano et al, 2012. The issues were then blocked in TNB buffer for 1 hour prior to incubation anti-DIG-POD (1:1000) for 18 hours at 4°C. Post-primary antibody incubation, tissues were washed using 0.1% PBT. For signal detection and amplification Alexa Fluor™ 594 Tyramide Reagent was used. Tyramide amplification solution was prepared as mentioned in the user guide. Tissues were incubated in TSA working solution for 8 minutes. Following this, an equal amount of Reaction stop reagent solution was added and further incubated for 1 minute. Post TSA reaction, tissues were PBS rinsed thrice for 5minutes and mounted in Vectashield.

## Acknowledgement

We thank B Lemaitre, N Silverman, I Ando, U Banerjee, S Cohen, P Ingham and M Crozatier and A. D. Shirras for reagents. Special thanks to Sushmit Ghosh and Kaustuv Ghosh for assisting with FISH and germ free experiments. We thank all members of the two labs for their valuable inputs. We thank IISER Mohali’s Confocal Facility, Bloomington Drosophila Stock Center, at Indiana University, VDRC (Vienna) and Developmental Studies Hybridoma Bank, University of Iowa for flies and antibodies. Models “Created with BioRender.com”. DBT Wellcome-Trust India Alliance Senior Fellowship [IA/S/17/1/503100] to LM and Institutional support to PR, NSD, SM and DBT-fellowship funding to AK for this study duly acknowledged.

## Inventory of Supplemental figures

Supplemental information contains 10 Supplemental Figures (Figure 1-figure supplement 1, Figure 2-figure supplement 1, Figure 3-figure supplement 1, Figure 4-figure supplement 1,Figure 4-figure supplement 2, Figure 5-figure supplement 1, Figure 5-figure supplement 2, Figure 6-figure supplement 1, Figure 6-figure supplement 2 and Figure 7 figure supplement 1.

**Figure 1 figure supplement 1:**
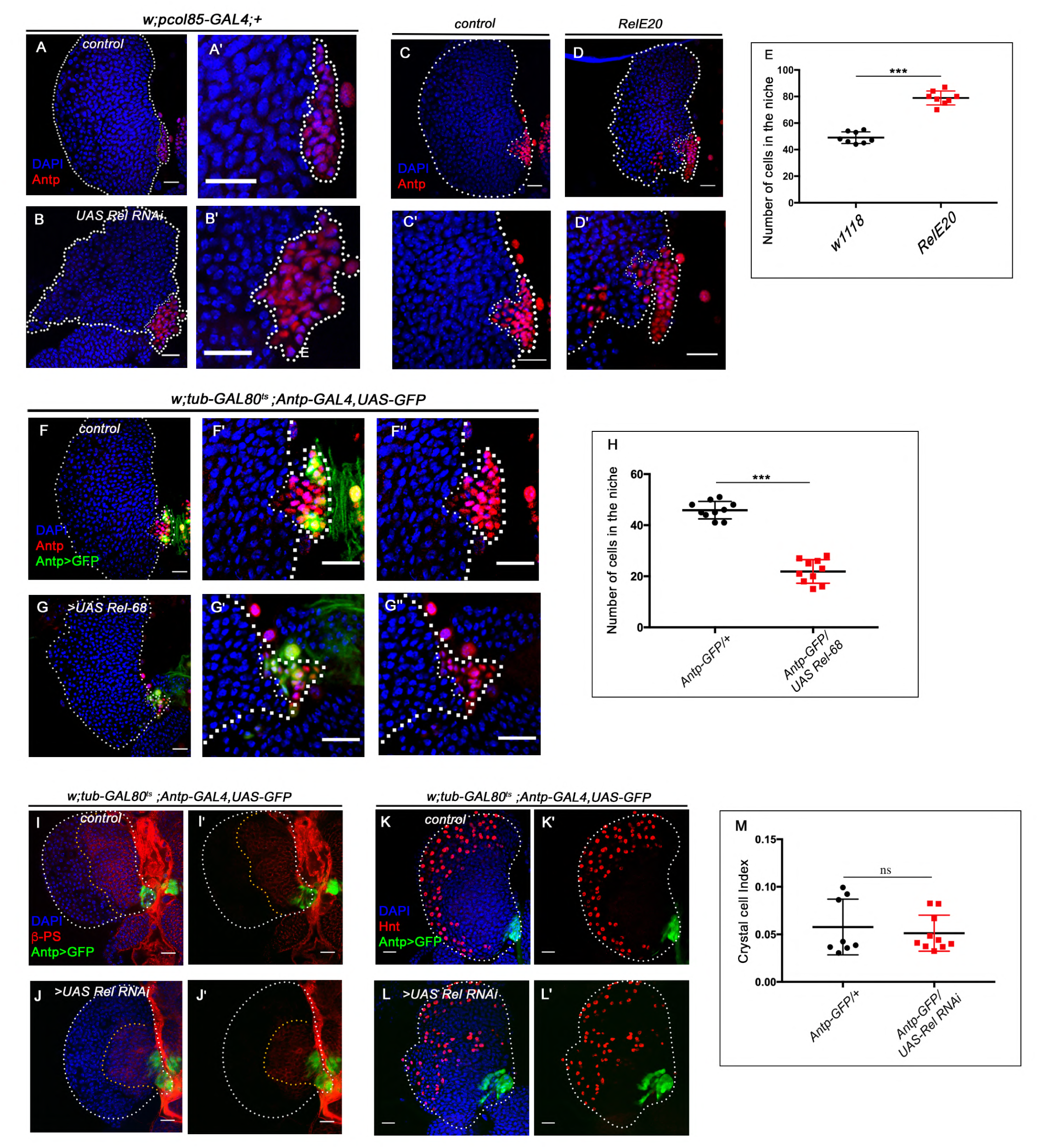
Relish negatively regulate niche cell proliferation. Genotypes of the larvae are mentioned in respective panels. Scale bar: 20µm **A-B’.** Effect of Relish loss from the niche using an independent GAL4 line, *pcol85-GAL4*. Compared to control (**A-A’**) down-regulation of Relish from the niche using *pcol85-GAL4* (**B-B’**) also leads to increased niche cell proliferation. **C-D’**. A substantial increase in niche number was observed in Relish mutant (*Rel^E20^*) (**D-D’)** when compared to control (**C-C’)**. (**E**) Quantitation of niche cell number in *Rel^E20^* mutant in comparison to control (n=8, P-value =9.03×10^-9^, two-tailed unpaired Student’s t-test). **F-G’’**. In comparison to control (**F-F’’)**, overexpression of Relish in the niche resulted in a reduction in niche cell number (**G-G’’)**. (**H**) Quantitation of niche cell number in Relish overexpression in comparison to control (n=10, P-value=3.3×10^-10^, two-tailed unpaired Student’s t-test). **I-J’-** Lamellocytes were not observed in Relish loss scenario (red, integrin β-PS-immunostaining. Loss of β-PS positive progenitor pool is further evident in Relish loss scenario compared to control (Compare **J-J’ to I-I’**) **K-L’**. In comparison to the control (**K-K’**), no significant change in crystal cell index (number of crystal cells/total number of cells in the lobe) was observed in Relish down-regulation scenario (**L-L’)**. (**M**). Quantitative analysis of crystal cell index in both control and Relish loss condition (n=8, P-value = 0.596, two-tailed unpaired Student’s t-test). The white dotted line mark whole of the lymph gland in all cases. The nuclei are marked with DAPI (Blue). In all panels, the age of the larvae is 96hrs AEH. Individual dots represent biological replicates. Error Bar: Standard Deviation (S.D). Data are mean±s.d. *P<0.05, **P<0.01 and ***P<0.001.

**Figure 2 figure supplement 1 :**
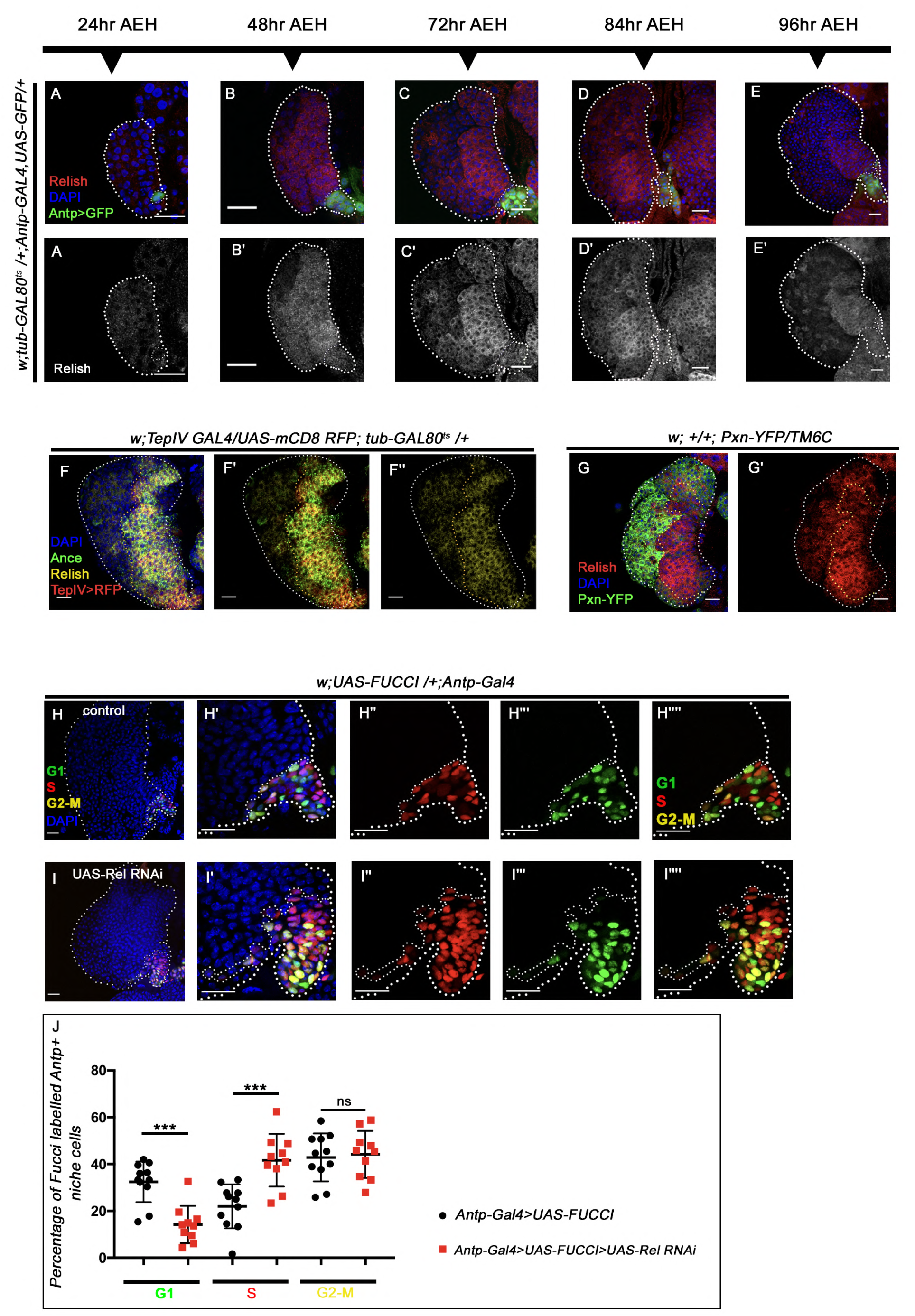
Relish expression starts beyond the second instar stage in the hematopoietic niche. The genotypes are mentioned in relevant panels. Scale bar: 20µm. **A-E’**. Expression of Relish (red, by antibody) at different developmental time points in the larval lymph gland (niche marked with *AntpGAL4>UASGFP)*. Observations were made at 24 hrs AEH (**A-A’**), 48 hrs AEH (**B-B’**), 72 hrs AEH (**C-C’**), 84 hrs AEH (**D-D’**), 96 hrs AEH (**E-E’**). Relish expression in the niche can be detected around 48 hrs AEH. **F-F’’**. Relish expression (yellow) in the progenitor cells co-localises with prohemocyte markers Ance (green) and TepIV (red). **G-G’.** Relish expression (red) is restricted to progenitor cells whereas it is downregulated in Pxn-YFP positive differentiated cells (green). **H-I’’’’**. Cell cycle status reported by Fly-FUCCI using niche-specific GAL4: *Antp-Gal4*. In control niche cells are mostly in G1 (green, **H’’’**), and G2-M (yellow, H**’’’’)** phase, while few are in S phase (red, **H’’**). Niche cells from where Relish function has been down-regulated were mostly in S, (red, **I’’**) and G2-M (yellow, **I’’’’**), and very less in G1 (green, **I’’’**) phase of the cell cycle. **J.** Quantitative analyses of the cell cycle status of control and Relish loss niches (n=10, P-value for G1=7.2×10^-5^, P-value for S=4.1×10^-4^, P-value for G2-M =.657), two-tailed unpaired Student’s t-test). The white dotted line mark whole of the lymph gland in all cases. In all panels age of the larvae is 96 hrs AEH, unless otherwise mentioned. The nuclei are marked with DAPI (Blue). Individual dots represent biological replicates. Error Bar: Standard Deviation (S.D). Data are mean±s.d. *P<0.05, **P<0.01 and ***P<0.001.

**Figure 3 figure supplement 1:**
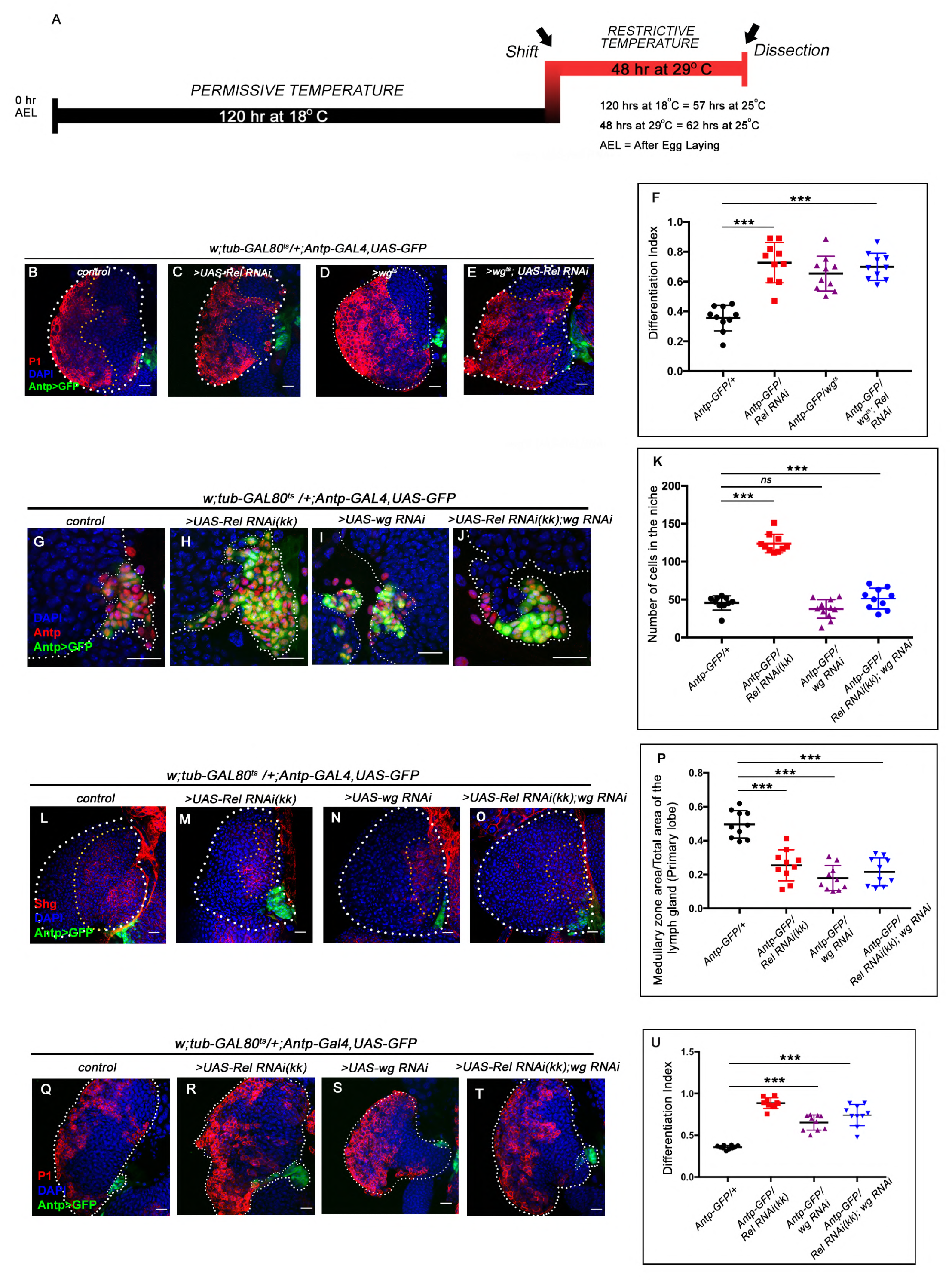
Downregulating wingless in Relish loss condition rescues niche cell proliferation but not differentiation. The genotypes are mentioned in relevant panels. Scale bar: 20µm. **A**. Scheme depicting the temperature regime followed for the rescue experiments (Figure 2D**-G*’,*** Figure 2I**-L and** Figure 2 **figure supplement 1 B-E)** for wingless mutant (*wg^ts^*). **B-E**. Increase in plasmatocyte population (marked by P1, red) was observed upon Relish (**C**) and in wingless downregulation (**D**) from the niche compared to the control (**B**). Simultaneous downregulation of wingless function in Relish loss genetic background did not rescue the increased differentiation (**E**). **F.** Statistical analysis of the data in (**B-E**) (n=10, P-value = 2.1×10^-6^ for control versus Relish RNAi, P-value = 6.80×10^−8^ for control versus wg^ts^; Relish RNAi; two-tailed unpaired Student’s t-test). **G-J**. The increased niche number observed upon Relish loss (**H**) is rescued upon reducing Wingless level by the wg-RNAi (**I**) in Relish loss genetic background **(J**). The rescued niche cell number is comparable to control (**G**). **K**. Statistical analysis of the data in (**G-J**) (n=10, P-value = 1.1×10^-11^ for control versus Relish RNAi^KK^, P-value = 3.15×10^−10^ for control versus wg RNAi; Relish RNAi^KK^; two-tailed unpaired Student’s t-test). **L-O**. Hematopoietic progenitors of larval lymph gland (red, reported by DE-Cadherin (Shg) immunostaining). Knocking down wingless function from the niche resulted in loss of Shg positive progenitors (**N**). Downregulating wingless using wg-RNAi in Relish loss genetic background was unable to restore the reduction in prohemocyte pool (**O**) observed in Relish loss (**M**) scenario in comparison to control (**L**). **P.** Statistical analysis of the data in (**L-O**) (n=10, P-value = 6.74×10^-6^ for control versus Relish RNAi^KK^, P-value = 4.03×10^−7^ for control versus wg RNAi; Relish RNAi^KK^; two-tailed unpaired Student’s t-test. **Q-T**. Increase in plasmatocyte population (marked by P1, red) was observed upon wingless (**S**) and Relish down regulation from the niche (**R**) compared with the control (**Q**). Simultaneous downregulation of wingless function in Relish loss genetic background did not rescue the increased differentiation (**T**). **U.** Statistical analysis of the data in (**Q-T**) (n=10, P-value = 2.9×10^-9^ for control versus Relish RNAi^KK^, P-value = 4.18×10^−5^ for control versus wg RNAi; Relish RNAi^KK^; two-tailed unpaired Student’s t-test. The white dotted line mark whole of the lymph gland in all cases. In all panels age of the larvae is 96 hrs AEH. The nuclei are marked with DAPI (Blue). Individual dots represent biological replicates. Error Bar: Standard Deviation (S.D). Data are mean±s.d. *P<0.05, **P<0.01 and ***P<0.001.

**Figure 4 figure supplement 1.**
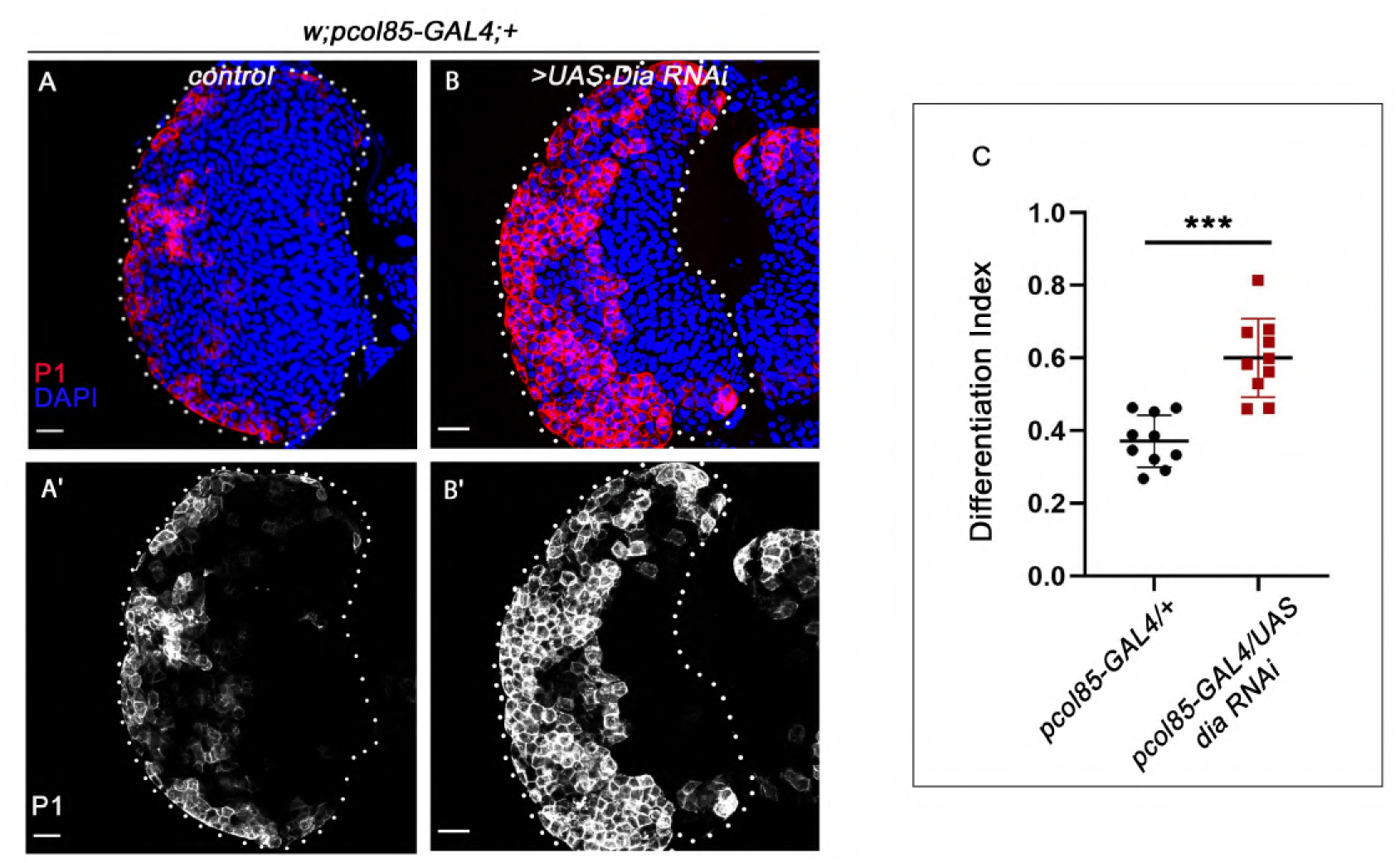
Loss of Diaphanous from the niche resulted in enhanced differentiation. Loss of Diaphanous (Dia), an actin polymerase from the niche caused ectopic differentiation of progenitors **(B-B’)** compared to control **(A-A’). C.** Differentiation index for Diaphanous loss niches compared to control (n=10, P-value= 4.28 x10^-5^; two tailed Students t-test). The white dotted line mark whole of the lymph gland in all cases. In all panels age of the larvae is 96 hrs AEH. The nuclei are marked with DAPI (Blue). Data are mean±s.d. *P<0.05, **P<0.01 and ***P<0.001.

**Figure 4 figure supplement 2 :**
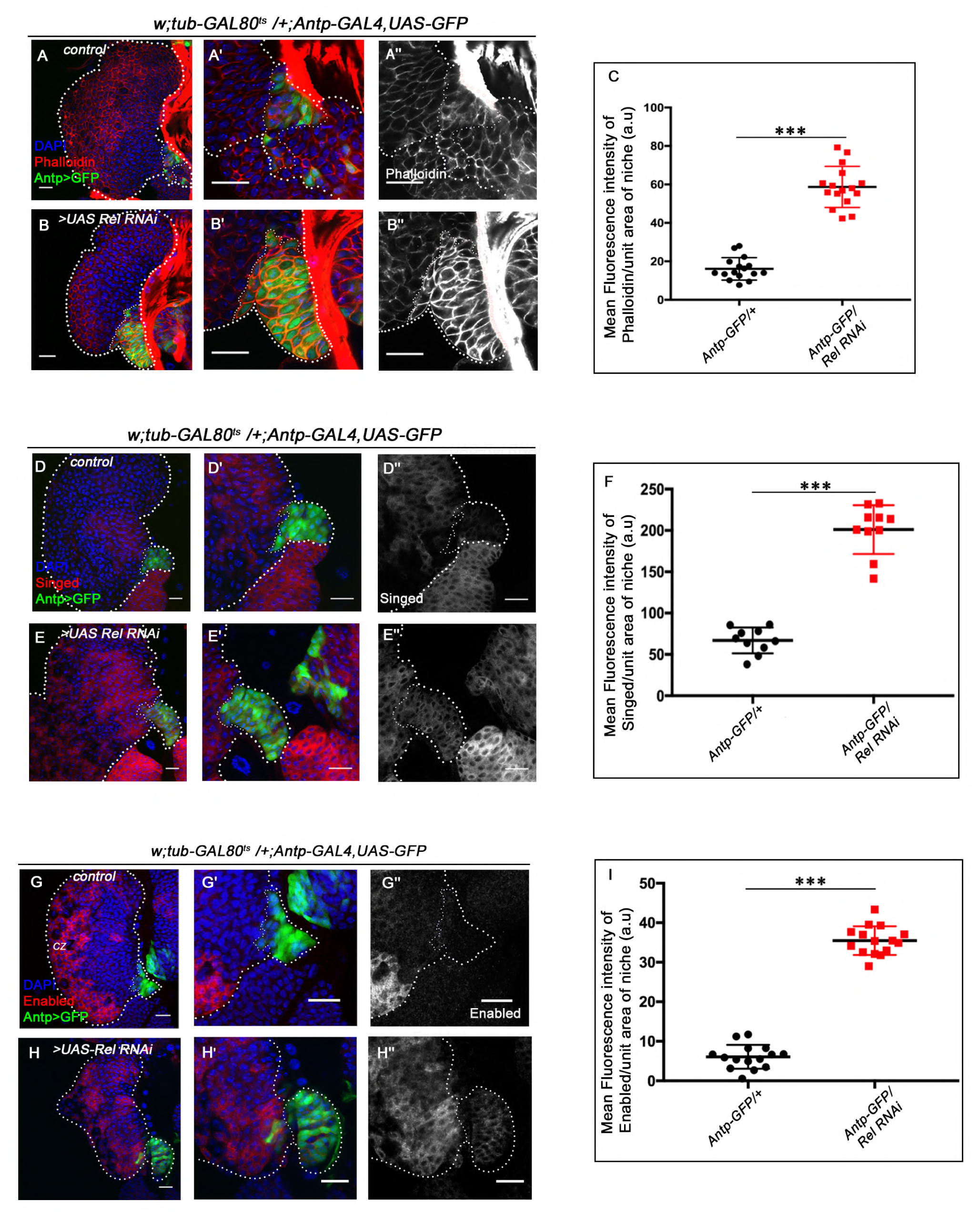
Loss of Relish from the niche resulted in upregulation of actin re-modellers. The genotypes are mentioned in relevant panels. Scale bar: 20µm. **A-B’’**. F-actin (visualised by Phalloidin, red) highly enriched in the plasma membrane of niche cells where Relish function is down-regulated (**B-B’’**) in comparison to that of control (**A-A’’**). **C.** Statistical analysis of fluorescence intensity showed significant increase in F-actin in Relish loss niches compared to control (n=10, P-value= 5.6×10^-9^, two tailed Students t-test). **D-E’’.** Expression of Singed, an actin bundling protein, is significantly upregulated in Relish loss niches (**E-E*’’***) compared to control (**D-D’’**). **F**. Statistical analysis of fluorescence intensity showed significant increase in Singed expression in Relish loss niches compared to control (n=15, P-value = 7.0 x10^-13^, two tailed Students t-test). **G-H’’**. Enabled an actin polymerase, which is normally absent from the niche cells of control (**G-G’’**) is unregulated upon Relish down regulation (**H-H’’**). (**I)** Statistical analysis of fluorescence intensity showed significant increase in Ena expression in Relish loss niches compared to control (n=15, P-value= 8.1 x10^-20^, two tailed Students t-test). The white dotted line mark whole of the lymph gland in all cases. In all panels age of the larvae is 96 hrs AEH. The nuclei are marked with DAPI (Blue). Individual dots represent biological replicates. Error Bar: Standard Deviation (S.D). Data are mean±s.d. *P<0.05, **P<0.01 and ***P<0.001.

**Figure 5 figure supplement 1:**
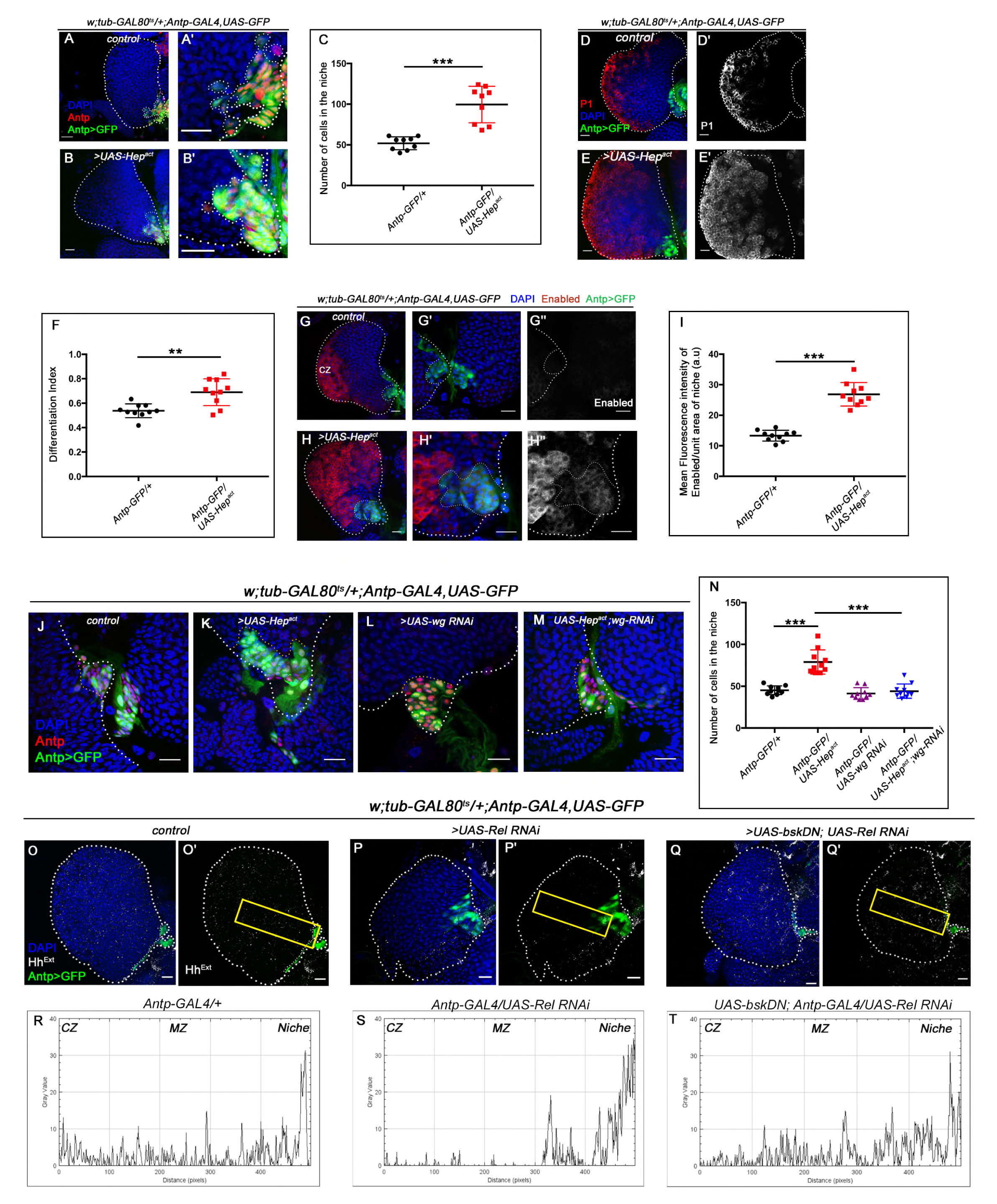
Ectopic activation of JNK signalling in the niche affects niche cell proliferation and progenitor maintenance. The genotypes are mentioned in relevant panels. Scale bar: 20µm. **A-B’**. An increase in niche cell numbers observed upon up-regulating JNK signaling using Hep^act^ in the niche (**B-B’**) compared to control (**A-A’**). **C.** Statistical analysis of the data in **A-B’** (n=10; P-value = 2.2×10^−4^ for control versus Hep^act^, two-tailed unpaired Student’s t-test). **D-E’.** A significant increase in differentiation observed JNK overexpression using Hep^act^ in the niche (**E-E’**) compared to control (**D-D’**). **F.** Statistical analysis of the data in **D-E’** (n=10, P-value = 1.7×10^−3^ for control versus Hep^act^, two-tailed unpaired Student’s t-test.) **G-H’’.** Robust increase in Enabled expression is observed when in Hep^act^ (**H-H’’**) compared to control (**G-G’’**). (**I**) Statistical analysis of the data in **G-H’’** (n=10; P-value = 2.1×10^−7^ for control versus Hep^act^, two-tailed unpaired Student’s t-test). **J-M**. Increase in niche cell numbers observed upon over-expressing Hep in the niche (**K**) is rescued to control levels (**J**) in a simultaneous loss of both Hep and wingless function from the niche (**M**). Loss of wingless using wg-RNAi had milder effect on niche cell number compared to control (compare **L** and **J**). **N**. Statistical analysis of the data in **J-M** (n=10; P-value = 2.20×10^−5^ for control versus Hep^act^, P-value = 1.08×10^−5^ for Hep^act^ versus Hep^act^; wg RNAi, two-tailed unpaired Student’s t-test). **O-Q’**. Reduced Extracellular Hh observed in the progenitors (Hh^Ext^) of Relish loss of function condition (**P-P’**) in comparison to those of control (**O-O’**), is significantly rescued in simultaneous loss of both Relish and JNK from the niche (**Q-Q’**). The yellow box in **O’, P’** and **Q’** denotes the area quantified in R, S and T respectively. **R-T.** The intensity profile of Hh^Extra^ in progenitors (along the rectangle drawn from PSC to Cortical zone housing differentiated cells in Figure 5 **figure supplement 1O’**, **P’** and **Q’**) reflects a stark decline in the level of Hh^Extr^ in Relish loss scenario **(S)** compared to control **(R)** which is rescued upon simultaneous loss of both Relish and JNK from the niche **(T)**. The white dotted line mark whole of the lymph gland in all cases. In all panels age of the larvae is 96 hrs AEH. The nuclei are marked with DAPI (Blue). Individual dots represent biological replicates. Error Bar: S.D. Data are mean±s.d. *P<0.05, **P<0.005 and ***P<0.0005.

**Figure 5 figure supplement 2 :**
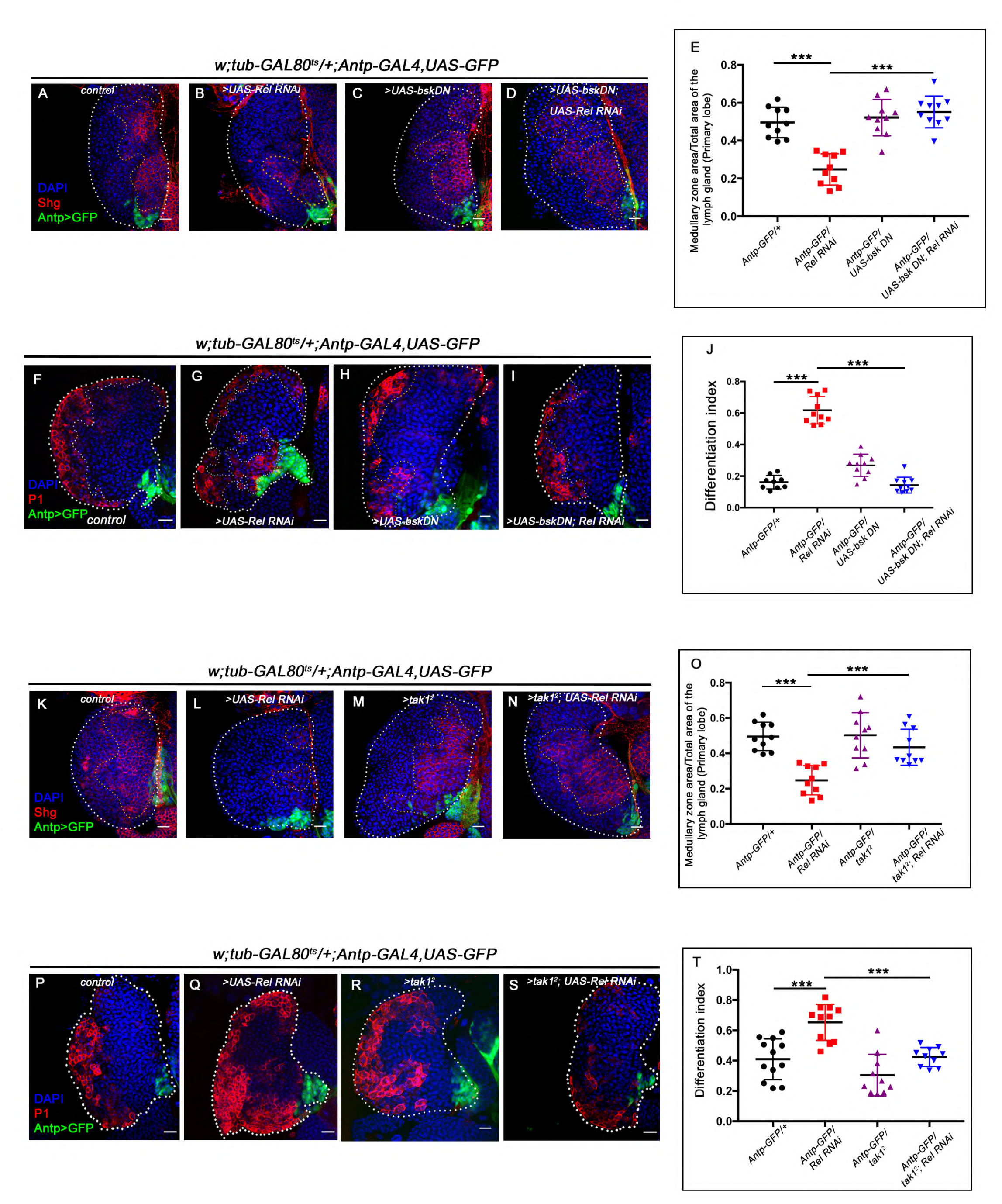
Downregulating JNK in Relish loss genetic background rescues progenitor loss and precocious differentiation. **A-D**. Hematopoietic progenitors of larval lymph gland (red, reported by DE-Cadherin (Shg) immunostaining). Knocking down JNK function from the niche did not have any effect on Shg positive progenitors (**C**). Downregulating bsk function in Relish loss genetic background was able to restore the reduction in prohemocyte pool (**D**) observed in Relish loss (**B**) scenario in comparison to control (**A**). **E**. Statistical analysis of the data in (**A-D**) (n=10, P-value = 2.26 ×10^−6^ for control versus Relish RNAi, P-value = 1.94×10^−7^ for bsk^DN^; Relish RNAi versus Relish RNAi; two-tailed unpaired Student’s t-test) **F-I**. Differentiation defect observed in Relish loss (**G**) was reverted to control (**F**) in a simultaneous knockdown of both Relish and JNK (**I**) from the niche. Loss of JNK from the niche had no significant effect on progenitor differentiation **(H)**. **J**. Statistical analysis of the data in **F-I** (n = 10, P-value = 1.5×10^−9^ for control versus Relish RNAi, P-value = 1.79×10^−8^ for bskDN; Relish RNAi versus Relish RNAi; two-tailed unpaired Student’s t-test). **K-N**. Loss of tak1 function from the niche did not have any effect on DE-Cad (Shg) positive progenitors (**M**). Downregulating tak1 in Relish loss genetic background could restore the reduction in prohemocyte pool (**N**) observed in Relish loss (**L**) scenario in comparison to control (**K**). **O**. Statistical analysis of the data in **K-N** (n = 10, P-value = 2.26×10^−6^ for control versus Relish RNAi, P-value = 3.1×10^−4^ for tak1^2^; Relish RNAi versus Relish RNAi; two-tailed unpaired Student’s t-test). **P-S**. Differentiation defects observed in Relish loss (**Q**) was reverted to control (**P**) in simultaneous loss of both Relish and tak1 function (**S**) from the niche. No significant change in differentiation was observed in tak1 loss from the niche (**R**). **T.** Statistical analysis of the data in **P-S** (n=10; P value = 1.5×10^−4^ for control versus Relish RNAi, P-value = 4.7×10^−5^ for; Relish RNAi versus tak1^2^; Relish RNAi; two-tailed unpaired Student’s t-test). The white dotted line mark whole of the lymph gland in all cases. In all panels age of the larvae is 96 hrs AEH. The nuclei are marked with DAPI (Blue). Individual dots represent biological replicates. Error Bar: Standard Deviation (S.D). Data are mean±s.d. *P<0.05, **P<0.01 and ***P<0.001.

**Figure 6 figure supplement 1:**
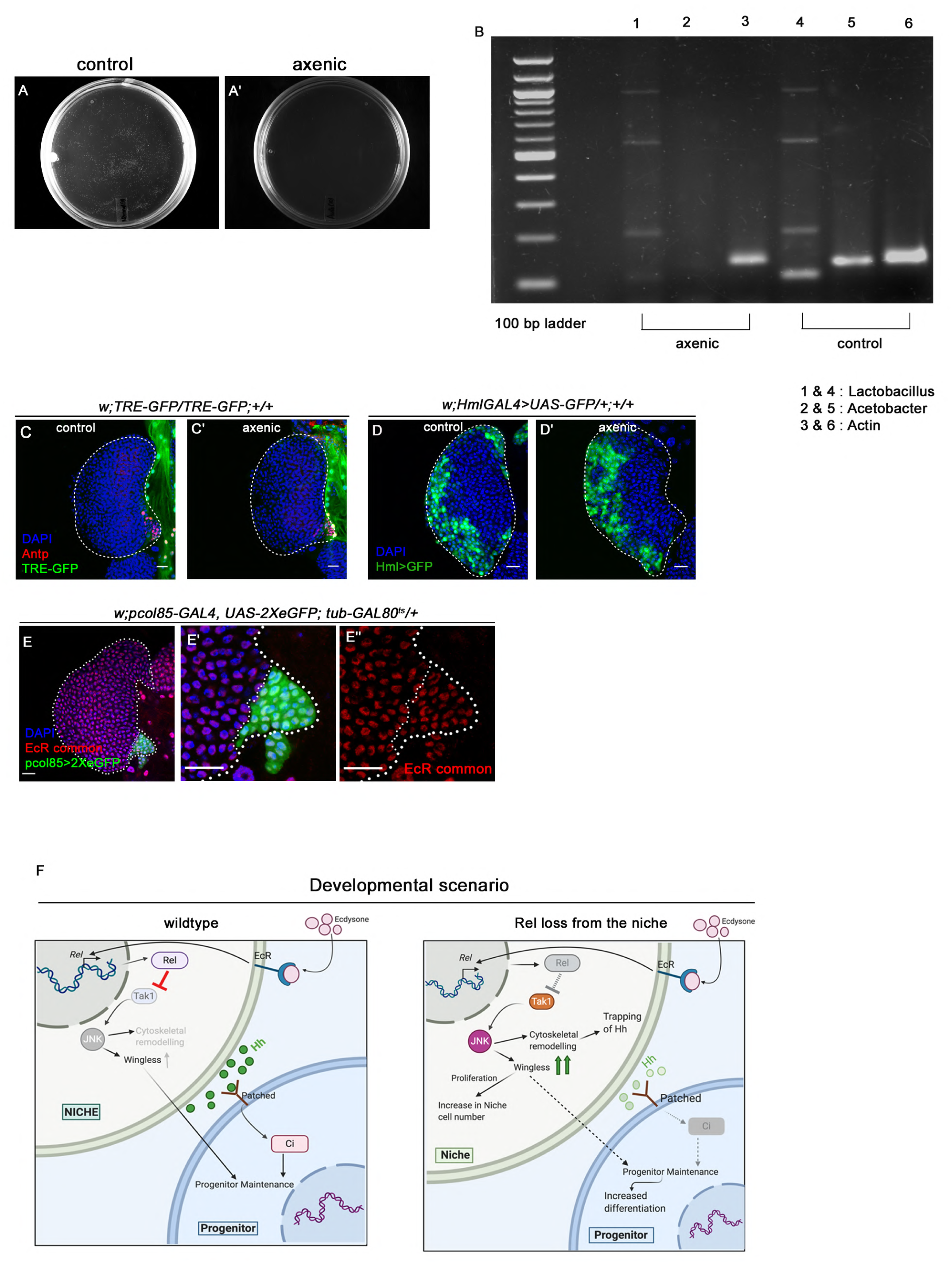
Ecdysone signaling is active in the hematopoietic niche. Genotypes of the larvae are mentioned in respective panels. Scale bar: 20µm **A-A’**. Larval homogenates were spread on LB Agar plates to check the presence of commensal gut microbiota. In control scenario (**A**) bacterial colonies were visible post incubation whereas in axenic condition no growth was observed on the plates (**A’**). **B**. The efficacy of removal of gut microflora was further checked by performing PCR analysis on DNA isolated from larval guts using 16S rDNA primers. *Drosophila* actin was used as control. Significant reduction in the amount of both *Lactobacillus* (compare lane 4 (axenic) with 1 (control) and *Acetobacter* (compare lane 5 (axenic) with 2 (control) species was observed in axenic condition compared to control scenario (compare lane 3 (axenic) and 6 (control). **C-C’**. TRE-GFP expression in the hematopoietic niche (visualised by Antp, red) in axenic condition (**C’**) is comparable to that of control (**C**). **D-D’.** Differentiation status (visualised by *Hml>GFP*, pan plasmatocyte marker) in axenic condition (**D’**) is comparable to control (**D**). **E-E’’**. Nuclear expression of Ecdysone receptor (red, EcR common) in the hematopoietic niche (green). **F**. Model depicting the developmental role of Relish in hematopoietic niche maintenance. Downregulation of Relish affects the proliferation and primary function of the niche by upregulated JNK signaling. Upregulated JNK disturbs niche homeostasis, thereby affecting progenitor maintenance. The white dotted line mark whole of the lymph gland in all cases. In all panels age of the larvae is 96 hrs AEH. The nuclei are marked with DAPI (Blue).

**Figure 6 figure supplement 2:**
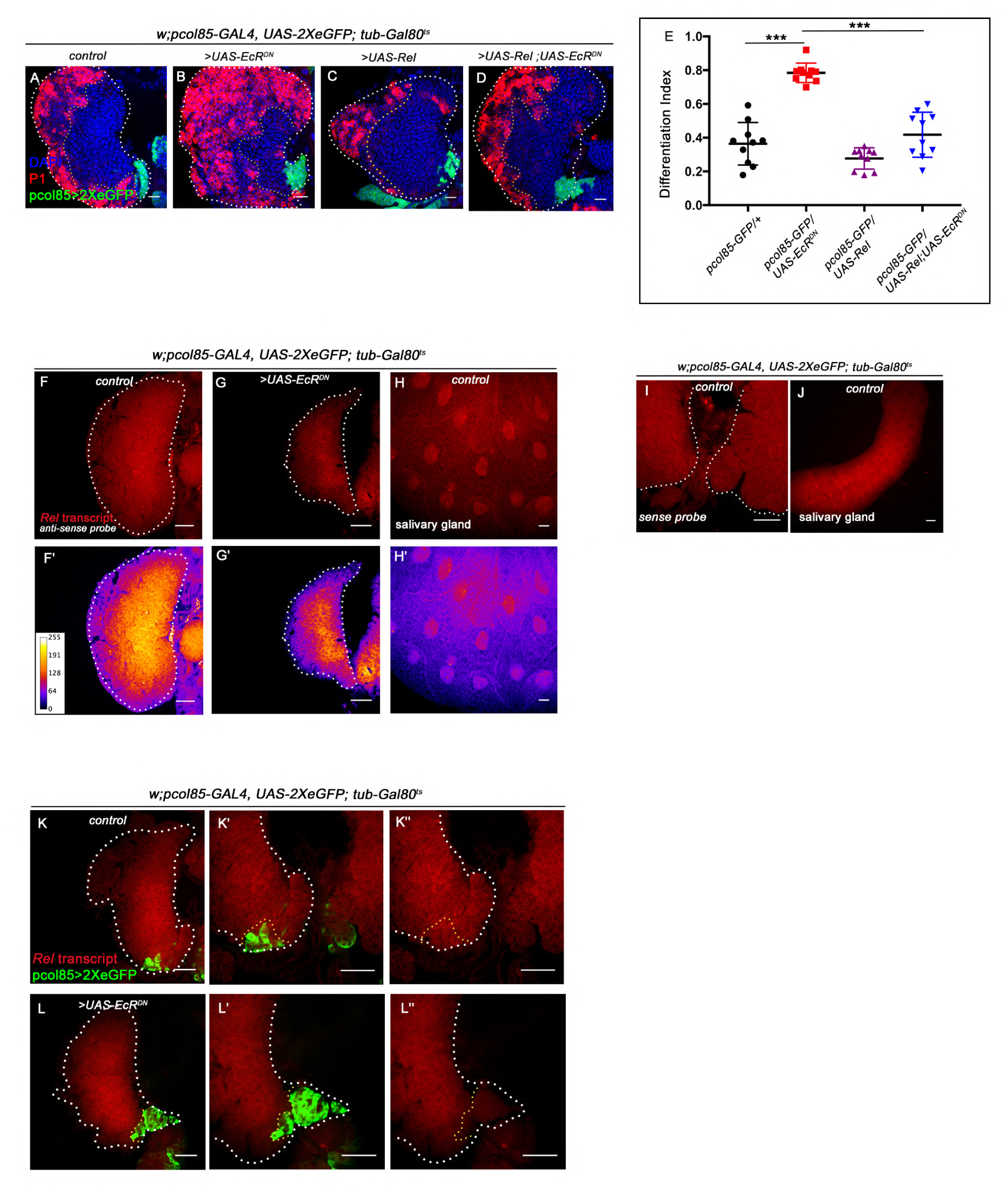
Relish expression is transcriptionally regulated by Ecdysone signaling in the hematopoietic niche. Genotypes of the larvae are mentioned in respective panels. Scale bar: 20µm **A-D**. Differentiation defects observed in EcR loss (**B**) was reverted to control (**A**) when Relish was overexpressed in EcR loss genetic background (**D**). Slight decrease in differentiation of progenitors were observed Relish overexpression scenario in the niche (Compare **A** and **C**). **E**. Statistical analysis of the data in **A-D** (n=10; P= 3.8×10^−7^ for control versus EcRDN, P= 3.3×10^−6^ for EcrDN versus UAS Rel; EcRDN, two-tailed unpaired Student’s t-test). **F-H’.** *Fluorescent in-situ hybridization* (FISH) analysis showing the expression of *Rel* transcript in the lymph gland of the control larvae (**F-F’**). Loss of EcR from the niche resulted in loss of Rel positive progenitors (**G-G’***). Rel* transcripts were also detected in salivary gland of the control larvae (**H-H’**). **I-J.** Sense probe showing nonspecific background expression in the control lymph gland (**I**) and salivary gland (**J**). **K-L’’.** Whole mount immunofluorescence (IF) and fluorescent *in-situ* hybridization (FISH) on third instar lymph gland. Compared to control (**K-K’’**) drastic reduction of the *Rel* transcript was observed in the niche from where EcR expression was downregulated (**L-L’’**). Please note the smaller size of the LG in G-G**’** reflects the peeling off of the cortical zone due to excessive differentiation around 96hr AEH in EcR loss from the niche. The increased differentiation renders fragility to the LG, which is unable to withstand harsh *insitu* process. The white dotted line mark whole of the lymph gland in all cases. In all panels age of the larvae is 96 hrs AEH. The nuclei are marked with DAPI (Blue). Individual dots represent biological replicates. Error Bar: Standard Deviation (S.D). Data are mean±s.d. *P<0.05, **P<0.01 and ***P<0.001.

**Figure 7 figure supplement 1:**
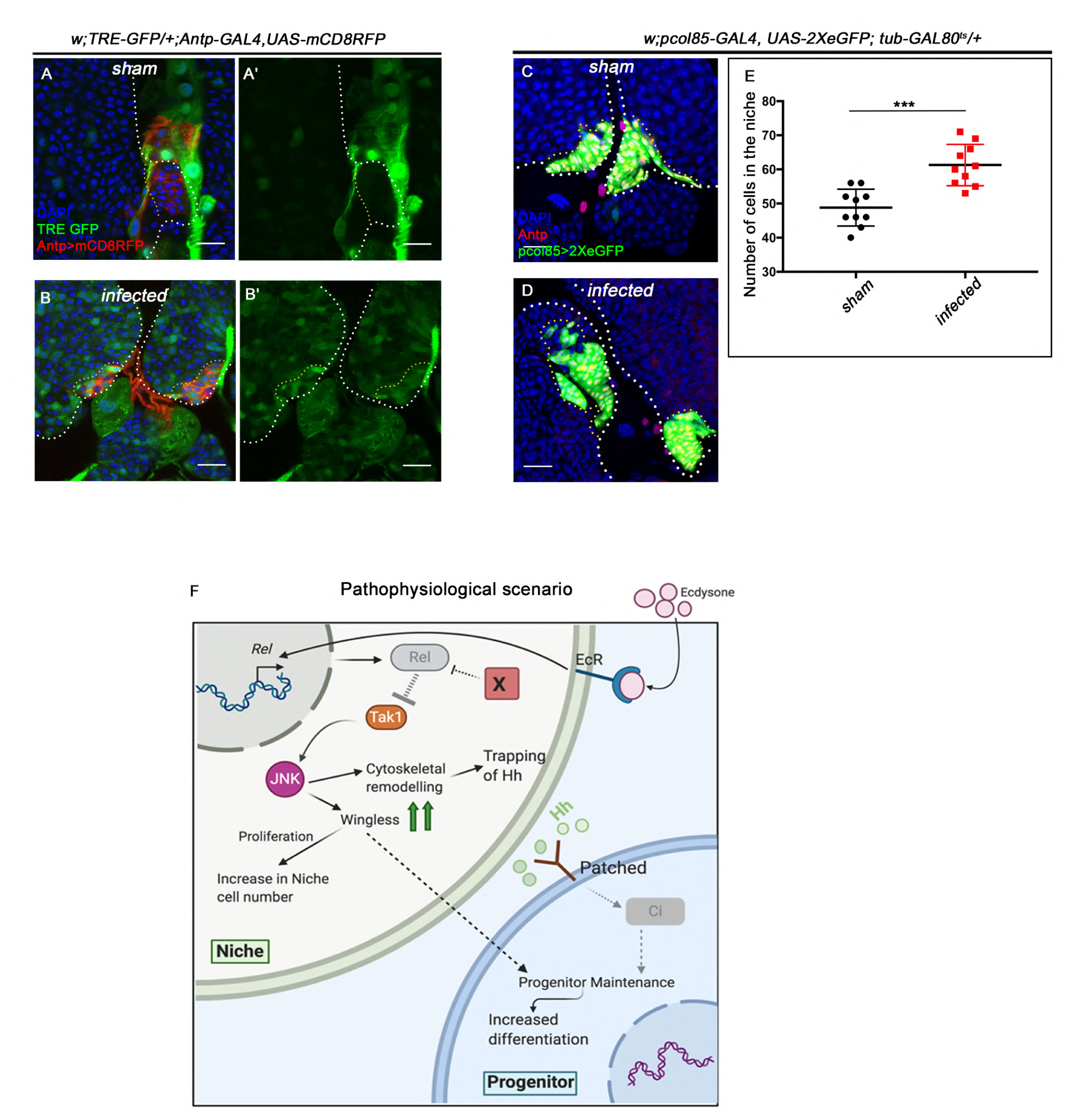
Upregulation in JNK signaling and increased niche cell proliferation was observed in the niche cells during infection. The genotypes are mentioned in relevant panels. Scale bar: 20µm. **A-B’.** An overall up regulation in JNK signalling (visualized by its reporter TRE-GFP (green) was observed in infected lymph glands (**B-B’**) compared to sham (**A-A’**). **C-D.** Significant increase in niche proliferation was observed in infected lymph gland niches **(D)** compared to sham infected **(C).** **E.** Statistical analysis of the data in **C-D** (n=10; P-value = 1.1×10^−4^ for sham versus infected, two-tailed unpaired Student’s t-test). **F.** Model based on current results depicting how upon bacterial challenge Relish expression is differentially modulated in the niche to bolster the cellular immune response by eliciting precocious differentiation of the lymph gland hemocytes. The white dotted line mark whole of the lymph gland in all cases. In all panels age of the larvae is 96 hrs AEH. The nuclei are marked with DAPI (Blue). Individual dots represent biological replicates. Error Bar: Standard Deviation (S.D). Data are mean±s.d. *P<0.05, **P<0.01 and ***P<0.001.

## Source Data Legend

**Source data 1**: Contains numerical data plotted in Figure 1C-Dʹ, Figure 1F-Gʹʹ, Figure 1H-Iʹ, Figure 1N-Oʹ and Figure 1 figure supplement 1C-Dʹ, Figure 1 figure supplement 1F-Gʹʹ and Figure 1 figure supplement 1K-Lʹ.

**Source data 2** : Contains numerical data plotted in Figure 2A-Bʹʹ, Figure 2C-Dʹʹ, Figure 2E-Fʹʹ, Figure 2G-Hʹʹ, Figure 2J-Kʹʹ and Figure 2 figure supplement 1H-Iʹʹʹʹ.

**Source data 3** : Contains numerical data plotted in Figure 3A-Bʹʹ, Figure 3D-Gʹ, Figure 3I-L and Figure 3 figure supplement 1B-E and Figure 3 figure supplement 1G-J, Figure 3 figure supplement 1L-O and Figure 3 figure supplement 1Q-T

**Source data 4** : Contains numerical data plotted in Figure 4A-Bʹʹ, Figure 4D-Eʹʹ, Figure 4G-Iʹ, Figure 4 figure supplement 1A-Bʹ, Figure 4 figure supplement 2A-Bʹʹ, Figure 4 figure supplement 2D-Eʹʹ and Figure 4 figure supplement 2G-Hʹʹ

**Source data 5** : Contains numerical data plotted in Figure 5A-Bʹ, Figure 5D-Gʹ, Figure 5I-L, Figure 5S-V and Figure 5 figure supplement 1A-Bʹ, Figure 5 figure supplement 1D-Eʹ, Figure 5 figure supplement 1G-Hʹʹ, Figure 5 figure supplement 1J-M, Figure 5 figure supplement 1O-Qʹ, Figure 5 figure supplement 2A-D, Figure 5 figure supplement 2F-I, Figure 5 figure supplement 2K-N and Figure 5 figure supplement 2P-S.

**Source data 6** : Contains numerical data plotted in Figure 6A-Cʹ, Figure 6E-Gʹ, Figure 6I-Kʹ, and Figure 6M-Oʹ, Figure 6Q-Tʹ, Figure 6 figure supplement 2A-D

**Source data 7** : Contains numerical data plotted in Figure 7A-Cʹ, Figure 7I-J, Figure 7L-M.

